# Mechano-Cas12a Assisted Tension Sensor (MCATS) for Massively Amplified Cell Traction Force Measurements

**DOI:** 10.1101/2022.10.26.513907

**Authors:** Yuxin Duan, Fania Szlam, Yuesong Hu, Wenchun Chen, Renhao Li, Yonggang Ke, Roman Sniecinski, Khalid Salaita

## Abstract

Cells transmit piconewton forces to mediate essential biological processes such as coagulation. One challenge is that cell-generated forces are infrequent, transient, and difficult to detect. Here, we report the development of Mechano-Cas12a Assisted Tension Sensor (MCATS) that utilizes CRISPR-Cas12a to transduce and amplify the molecular forces generated by cells. We demonstrate the power of MCATS by detecting the forces generated by as few as ~10^3^ human platelets in a high-throughput manner. Platelet forces are significantly inhibited when blood samples are treated with FDA-approved drugs such as aspirin, eptifibatide(integrilin®), 7E3(Reopro®), and ticagrelor (Brelinta®). Because MCATS requires <5uL of blood/measurement, a single blood draw can generate a personalized dose-response curve and IC_50_ for this panel of drugs. Platelet activity and force-generation are tightly associated, and hence MCATS was used to quantify platelet dysfunction following cardiopulmonary bypass (CPB) in a pilot study of 7 cardiac patients. We found that MCATS detected platelet dysfunction which strongly correlated with the need for platelet transfusion to limit bleeding. These results indicate MCATS may be a useful assay for clinical applications.

---

The ability for cells to generate mechanical forces is central to a wide range of biological processes ranging from immunology to coagulation and plays an important role in numerous pathologies such as cancer.^1–4^ Therefore, developing methods to quantify cell-generated force has important biomedical applications. A central challenge in this field is that molecular forces that are sensed and transduced by cells are fairly weak, at the scale of piconewtons (pN)^5^ and are highly transient and infrequent ^6^. Our lab and others recently developed DNA-based tension sensors that respond to cell generated pN forces and can be imaged using high-power microscopes with single molecule sensitivity.^7^ Despite the advances in mechanobiology enabled by DNA tension sensors,^3, 5, 7^ the use of these probes remains limited because of the weak signal and need for dedicated microscopy instrumentation. To facilitate the study of mechanobiology, it is important to develop facile, robust, and sensitive assays that can be broadly adopted by the community.

In biochemistry and molecular biology, weak or difficult to detect signal is typically enhanced by using catalytic amplification reactions such as PCR and ELISA. However, these assays have poor compatibility with live cell measurements. For example, thermal cycling in PCR would destroy most cells. An emerging class of enzymatically amplified reactions that are used in molecular diagnostics are based on clustered regularly interspaced short palindromic repeats (CRISPR) and CRISPR-associated proteins (Cas).^8, 9^ One notable Cas enzyme used in diagnostics is CRISPR-Cas12a (Cpf1) which is a class 2 type V-A enzyme that is loaded with single-stranded guide RNA (gRNA) and is activated upon binding to a complementary single stranded activator DNA. Upon activation of Cas12a, the enzyme undergoes a conformation change that unleashes its indiscriminate cleavage activity (trans activity) which hydrolyzes any ssDNA in proximity.^10^ The nuclease activity of Cas12a is robust, highly efficient with *k*_cat_/*K*_m_ ~ 10^6^-10^7^ M^−1^s^−1^, and thus has been used for a number of diagnostic assays for nucleic acid sensing^11^ such as DETECTR^12^ and for metal ion detection^12, 13^ Given the sensitivity and specificity of Cas12a based assays, we were inspired to adopt Cas12a enzyme to address the limited signal in cellular tension sensing assays.

Specifically, we developed a Mechano-Cas12a Assisted Tension Sensor (MCATS), which is an ultrasensitive fluorescence-based assay to detect the molecular forces generated by cells. In MCATS, the activator is a ssDNA anchored to a surface, such as a glass slide. The activator is concealed by hybridization to a complementary strand that is in turn conjugated to a peptide such as, cyclo-Arg-Gly-Asp-Phe-Lys (cRGDfK), or any protein ligand specific to the cell receptor of interest (**Figure 1a-b**). When cells are seeded on this surface, surface receptors such as integrins bind to the cRGDfK ligand on the duplex and apply forces. Forces that exceed the mechanical tolerance of the duplex lead to its rupture, exposing the activator (bottom strand) and thus triggering Cas12a nuclease activity. Upon activation, Cas12a will indiscriminately and catalytically cleave fluorogenic single stranded DNA reporter for amplification. Because Cas12a is highly efficient, its activation generates a massive fluorescence signal output that can be measured using a conventional fluorometer or plate reader for facile and high throughput readout.

**Figure 1.**
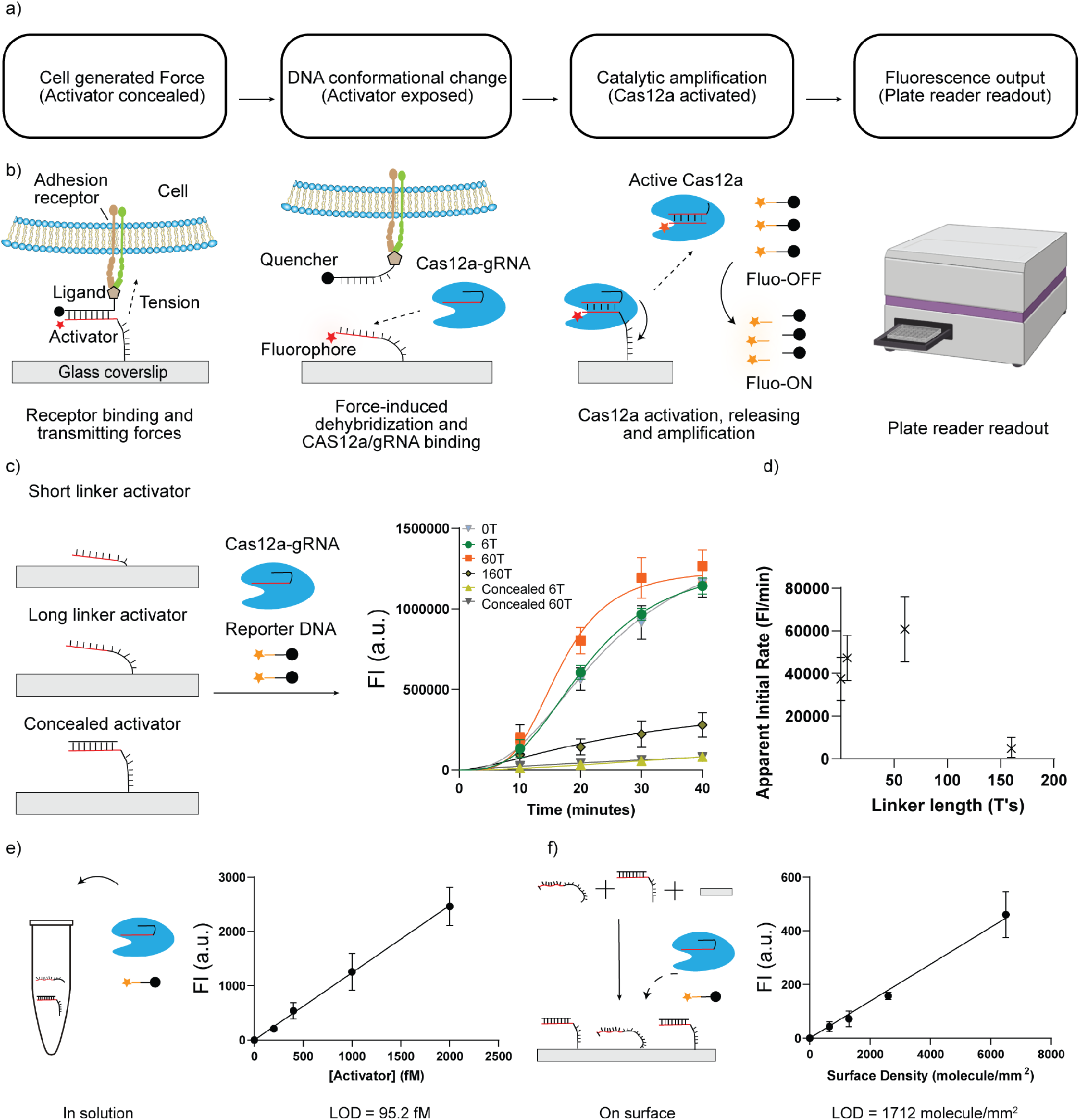
Scheme and characterization of MCATS. a-b) Flow chart and scheme depicting MCATS. Molecular traction forces mechanically melt the duplex probe and reveal an activator sequence that triggers Cas12a to cleave reporter strands in solution. The activator was tagged with Atto647N to aid in quantification and validation. c) Characterization of Cas12a activity for surface-tethered activators of 0, 6, 60, and 160 polyT linkers along with negative controls using concealed (duplexed) activators. Plot of time-dependent fluorescence intensity measured using a conventional plate reader. Error bars represent S.E.M. obtained from three independent experiments. d) Plot of Cas12a apparent initial cleavage rate as a function of linker length for immobilized activator. The apparent rate was determined using the change in fluorescence between t = 10 min and 20 min. e) Plot of fluorescence intensity at *t* = 1 hr as a function of activator concentration for reactions employing 20 nM Cas12a-gRNA complex and 100 nM reporter DNA. Each reaction also contained 100 nM of dsDNA to mimic the concealed activator. LOD = 3.3(standard deviation of blank/slope of calibration curve) = 95.2 fM. f) Plot of fluorescence intensity as a function of surface density of activator. Each reaction contained 20 nM Cas12a-gRNA complex and 100 nM reporter DNA. Surface displayed a binary mixture of activator and scrambled DNA. LOD = 1712 molecule/mm^2^.

MCATS is a platform technology and can be applied to study many different types of cells and different biological processes. As a proof-of-concept demonstration, we used MCATS to investigate the forces generated by human platelets because of the importance of mechanical forces in platelet function. Indeed, recent studies using micropatterned polymer structures showed that platelets forces can be used to detect underlying genetic clotting disorders^13^, and for predicting trauma-induced coagulopathy. ^14 15^ Because MCATS only requires ~5 uL of blood or less to conduct each measurement, a typical blood draw (~5 mL) allows one to run ~1000 assays in a rapid manner. We leveraged this capability to screen the activity of a panel of clinically approved anti-platelet drugs such as aspirin, integrilin, abciximab, and Brelinta. These experiments demonstrate the potential of MCATS for personalized tailoring of anti-coagulant drugs and may help guide therapeutic intervention in the clinic.^16^ We also applied MCATS to detect platelet dysfunction following cardiopulmonary bypass (CPB). In a pilot study of seven CPB patients, we found that the change in MCATS signal strongly correlated with the need for platelet transfusion. More broadly, MCATS offers an accurate, rapid, and cost-efficient method to detect molecular forces generated by cells and will hence open the door to integrating mechanical measurements in the clinic.

## Results

### Design and optimization of MCATS

We first designed a double stranded DNA tension probe that can be mechanically ruptured by cell generated forces to expose the immobilized activator. Because the DNA tension probe is specifically designed without PAM sequence, the concealed double strand tension probe fails to activate the Cas12a nuclease without mechanical activation. Upon mechanical denaturation of the duplex, the exposed activator can then activate gRNA/Cas12a complex to cleave single strand reporter DNA. We intentionally designed the reporter DNA with a short poly T sequence to minimize secondary structure and then tagged its two termini with a quencher-fluorophore pair that dequenched upon DNA hydrolysis. Sequence of all oligonucleotides used in this work are provided in **Table S1.**

Given that surface-tethered Cas12a had not been reported in the literature, and the potential for hindered activity due to immobilization^17^, we first measured the kinetics of Cas12a when the activator is immobilized on a surface and compared it to reactions where the activator was in solution. In this assay, biotinylated activator (100 nM) was anchored to streptavidin-coated surfaces by incubating for 1 hr at RT. Based on our previous surface calibration, this procedure generates a DNA density of 1330 ± 60 molecules/μm^2^.^18^ Next, the gRNA-Cas12a complex (20 nM) and reporter DNA (100 nM) were added to the surface and a fluorescence plate reader was used to monitor the fluorescence signal in each well of the 96 well plate. The measurement showed that ss-activator triggered the Cas12a nuclease and generated a strong fluorescent response. In contrast, the ds-activator (concealed activator) only showed minimal signal (**Figure 1c**).

Next, we tested whether adding a spacer to the activator may boost Cas12a cleavage rates. We tested four activators with different length polyT spacers of 0, 6, 60, and 160 nt. The surface density calibration showed that 6 and 60 polyT spacers only reduced surface density slightly (~18%) while the long 160nt spacer caused the surface density to decrease to 60% of the 0-nt spacer probe (**Figure S1**). The apparent initial rate constant (fluorescence signal change between t =10 and 20 min) for these immobilized activators is plotted in **Figure 1d** and showed that the apparent initial rate constant was enhanced with longer spacers with the exception for the 160 nt activator. The enhanced activity with longer spacers is likely due to reduced steric hinderance, but this effect is offset by the reduced effective activator density at extreme spacer lengths. Notably, we expect that the Cas12a will release the activator from the surface with suitable length spacers. This is supported by the observation that activator surface density is diminished by 70% for the 60 polyT spacer after 1 hr of adding the Cas12a (**Figure S2**). All subsequent work with MCATS employed activator with 60T spacer due to its superior signal amplification.

To further optimize MCATS, we measured Cas12a activity as a function of assay temperature, buffer, and reaction time (**Figure S3**). The results indicate the Cas12a work best at 37 °C, in cell culture medium with 10mM of Mg^2+^. We also compared the signal to noise ratio (S/N) of the assay using two reporter oligonucleotides (**Figure S4**). The reporter strand tagged with Atto565N-BHQ2 shows a ~two times higher S/N compared with FAM-Lowa black reporter strand because of its better quenching efficiency. Using the optimized conditions, we investigated the limit of detection (LOD) of our assay both for surface tethered activator as well as soluble activator as a reference (**Figure 1e-f**). To tune the activator surface density, we created surfaces comprised of a binary mixture of single stranded DNA activator and the blocked (double stranded) control DNA. We maintained a total activator solution concentration of 100 nM but tuned the ratio between the blocked DNA and single stranded activator. We then added Cas12a-gRNA and fluorogenic reporter to the well and measured the final fluorescence intensity after 1hr of enzyme activity. The LOD was then inferred from the ratio of 3.3 × standard deviation of the background normalized by the slope of the fluorescence versus concentration plot. The results showed that the LOD for nucleic acid sensing was 95.2 fM in solution and 1712 molecule/mm^2^ on a surface. This analysis provides the basis for using MCATS to sensitively detect molecular forces generated by cells.

### Rapid, robust, ultrasensitive detection of cellular tension with MCATS

We next applied the MCATS assay to detect integrin-mediated cell traction forces in immortalized cell lines. Integrins are a family of heterodimeric cell surface adhesion receptors that bridge the cellular cytoskeleton with the extracellular matrix (ECM) to mediate a variety of processes including cell adhesion, and migration.^2^ Previous research has shown that integrin receptors can apply pN forces which are sufficient to mechanically denature DNA duplexes.^19^ Hence, measuring integrin receptor forces with NIH/3T3 cells is an appropriate model to validate the MCATS assay.

As is shown in **Figure 2a**, DNA duplexes that present cRGDfk ligand and concealed Cas12a activator were immobilized on the surface. The conjugation of ligand to duplex was achieved via copper(I)-catalyzed azidealkyne cycloaddition and was verified with ESI-MS (**Figure S5, Table S2**). NIH/3T3 fibroblast cells were then seeded on these surfaces for 1 hr.

**Figure 2.**
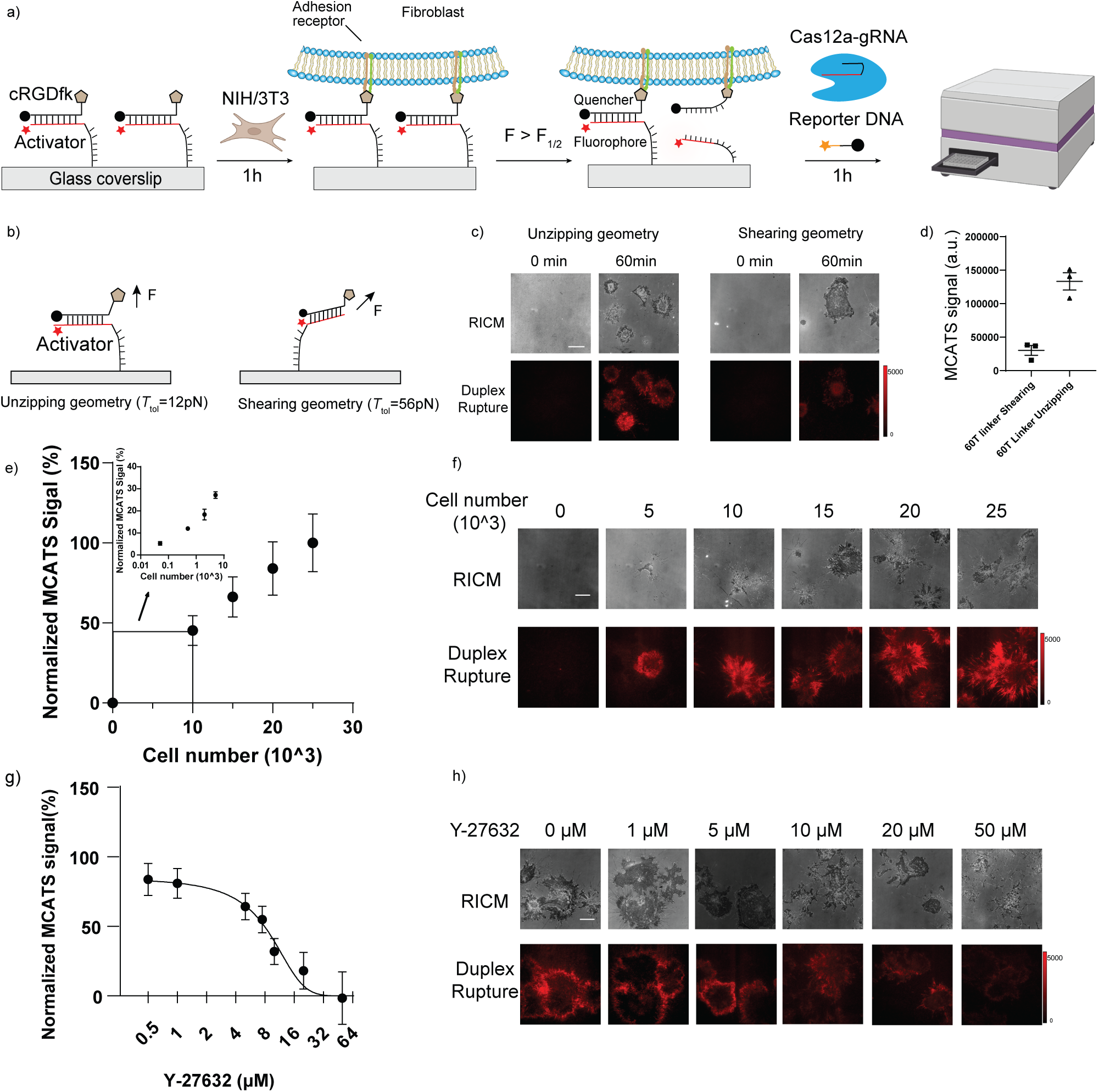
MCATS demonstration using fibroblasts. Schematic of MCATS assay to study NIH/3T3 integrin-mediated forces. b) Schematic comparing designs of concealed activator in shearing and unzipping geometries. c) Representative RICM and fluorescence images of cells cultured on unzipping (12 pN) and shearing (56 pN) surfaces at t = 0, and 1hr after cell seeding. Scale bar = 12 μm. The fluorescence emission is due to dequenching of Atto647N following top strand denaturation. The intensity bar indicates the absolute range of intensity values for each image. d) Plot shows MCATS signal measured using plate reader for cells incubated on T_tol_= 12 pN and T_tol_= 56 pN surfaces. Error bars represent S.E.M. from *n*=3 independent experiments. e) Plot showing plate reader measured MCATS signal as a function of the number of cells seeded. Error bar represents S.E.M. from three independent experiments. Inset shows signal in the regime of low numbers of cells (50-5000 cells). f) Representative RICM and duplex rupture (red) fluorescence images for unzipping probes. Scale bar = 12 μm. g) Plot of MCATS signal as a function of Y-27632 concentration. Drug was incubated for 30 min prior to seeding cells on the surface. Error bar represents S.E.M. from 3 independent experiments. Mechano-IC50 was calculated by fitting plot to a standard dose-response function: normalized signal = 100/(1+[drug]/IC50). The values were normalized to the signal obtained from the 25,000 cells/well samples without drug treatment. All measurements were background subtracted using negative control wells lacking cells. h) Representative RICM and duplex rupture (red) fluorescence images of drug-treated fibroblasts at t = 1 hr after seeding. Scale bar = 12 μm.

We designed two types of DNA duplexes that have identical sequence and thermal melting temperatures, but different geometries and mechanical tolerances (**Figure 2b**). When the activator is anchored through its 5’ terminus, the cRGDfK ligand is presented on the 3’ terminus of the top strand. Hence this probe denatures through an unzipping process which has a lower activation barrier and displays a mechanical rupture threshold of 12 pN. In contrast, when the activator is anchored through its 3’terminus, the probe denatures by shearing which has a larger mechanical threshold of 56 pN.

The same number of cells (25,000 cells) were incubated on the two types of surfaces for 1 hr before running the Cas12a amplification assay. As expected, we observed an increase in fluorescence signal for both the 12 pN and 56 pN probes (tagged with fluorophore quencher pairs) due to mechanical denaturation of the duplex (**Figure 2c**). As expected, the probes in the unzipping geometry (12 pN) were more significantly denatured compared to shearing mode probes (56 pN), in agreement with past literature (**Figure 2b-c**). We next added the Cas12a and reporter DNA to trigger the MCATS assay for 1 hr and then measured bulk fluorescence (λ_ex_ = 540 nm and λ_em_=590 nm) using the plate reader (**Figure 2d**). Importantly, the 12 pN unzipping mode probes also generated a greater MCATS signal, reflecting the greater density of exposed activator. MCATS produced over 100-fold greater signal compared to mechano-HCR which confirms that this assay is more sensitive and offers a simplified experimental process as no washing steps are required (**Figure S6**).^18^

We further validated the MCATS assay by seeding increasing numbers of NIH/3T3 cells in 96 well plates and measuring associated MCATS signal in each well. We observed that increasing cell densities led to greater MCATS signal (**Figure 2e-f**). Impressively, we found that MCATS can detect the tension generated from as few as 50 fibroblasts in a 96 well plate using a conventional plate reader (**Figure 2e inset**). We further tested whether MCATS can be used to produce a dose response curve for NIH-3T3 cell incubated with Rho kinase inhibitor, Y-27632, which targets the phosphorylation of myosin light chain and therefore dampens forces transmitted by focal adhesions. We pretreated NIH/3T3 cells with a range of Y-27632 concentrations (0-50 μM) for 30 min and then ran MCATS with the drug treated cell. MCATS signal showed a dose-dependent reduction as a function of increasing Y-27632 concentration (**Figure 2g-h**), indicating that plate-reader based MCATS readout can report cell forces modulated by MLC inhibition. By fitting the data to a standard dose-response inhibition function (signal =100/(1+[drug]/IC50)), we found that the mechano-IC_50_ = 7.9 μM (95% CI = 5.5 −11.6 μM), which matches previous literature reporting IC_50_ of 5-10 μM.^20^

### High-throughput determination of platelet inhibitors’ influence on platelet tension

We next investigated the MCATS signal of human platelets under the influence of different antiplatelet drugs. Platelets are primary cells and are well suited for analysis by because contractile forces are critical in platelet function in forming clots that mechanically resist blood shear flow and seal a wound. Anti-platelet agents are some of the most commonly prescribed drugs in the world. Currently, 30 million Americans take anti-platelet medications such as daily aspirin to reduce the risk of cardiovascular events, but ~16.6 of 100 patients experience bleeding as a side effect and 5% are resistant to aspirin.^21^

We tested multiple types of FDA-approved antiplatelet drugs: aspirin which inhibits the activity of cyclooxygenase (COX), the integrin αIIbβ3 antagonists eptifibatide, 7E3 (monoclonal antibody) as well as the P2Y12 inhibitor Ticagrelor.^22–24^ As is shown in **Figure 3a**, we first purified human platelets from blood samples collected in collection tubes containing sodium citrate or EDTA from donors. The sample was then centrifuged for 12 min at 140 *g* (with 0.02U Apyrase) to obtain platelet-rich plasma (PRP). Then platelets were prepared from PRP by centrifugation for 5 min at 700 *g* with 3 μM PGE-1. The platelets were then resuspended in Tyrodes buffer and centrifuged for 5 min at 700 *g* with 3 μM PGE-1. Finally, platelets were resuspended in Tyrodes buffer. It is worth noting that Apyrase is important in the platelet purification process to prevent hemolysis mediated platelet aggregation (**Figure S7d**). In additional controls, we found that tubes containing either EDTA or citrate will not influence platelet tension after purification. (**Figure S7d**)

**Figure 3.**
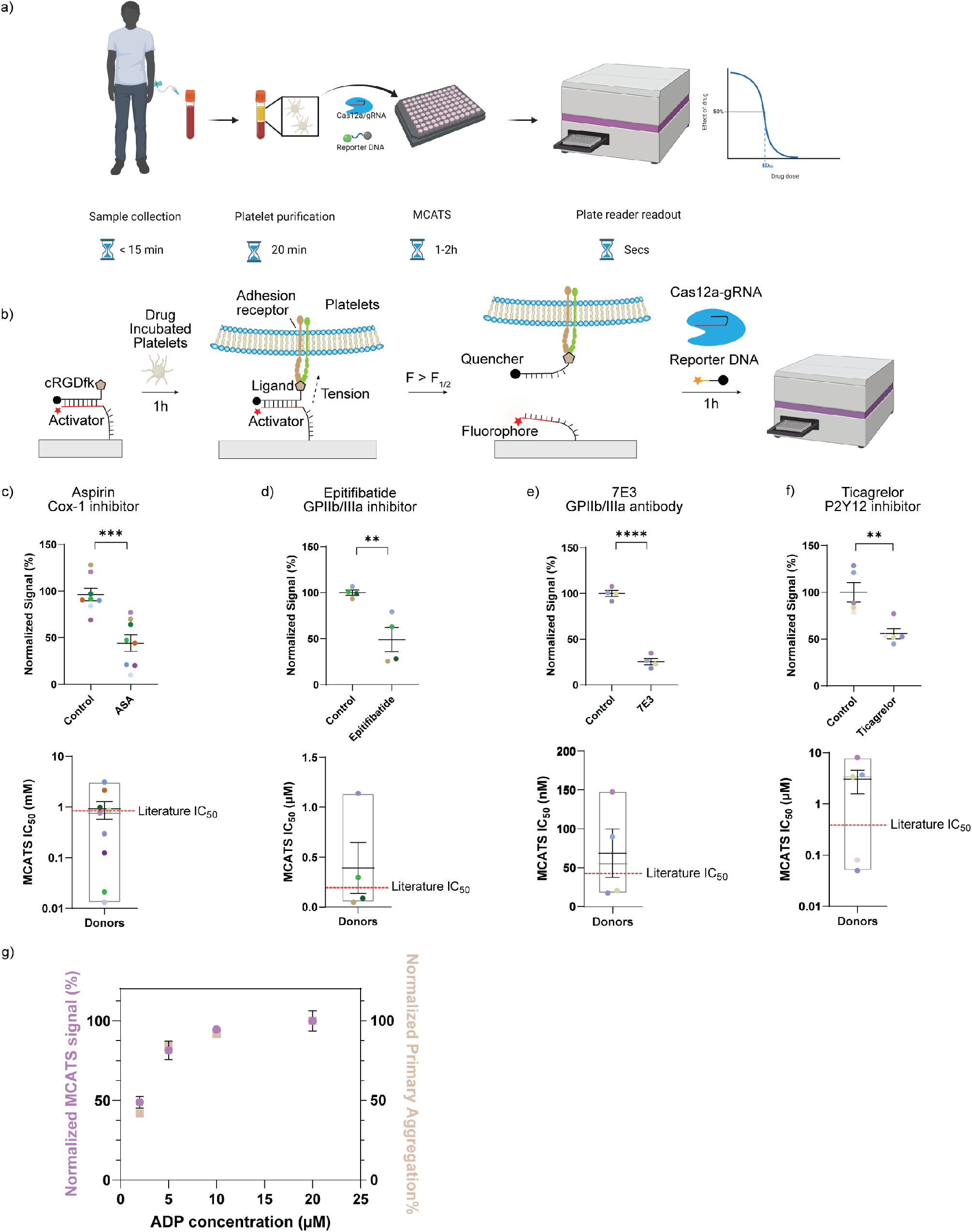
MCATS for personalized antiplatelet drug-sensitivity measurement. a) Schematic showing overall MCATS workflow using human platelets. b) Schematic showing mechanism of MCATS to quantify platelet-mediated forces. c-f) Plots of MCATS signal for platelets before and after inhibition using 0.1 mM aspirin, eptifibatide (1 μM), 7E3 (10 μg/ml), and Ticagrelor (1 μM). Donor platelets were treated with the inhibitor for 30 min at RT prior to performing MCATS. Each data point represents the average of n=2 or 3 measurements from a single donor. The mean and S.E.M. are shown for each drug treatment. Data was normalized to average of non-treated group. Significance is calculated with two tailed student t-test. The bottom set of plots show the mechano-IC_50_ calculated from a dose-response titration of six drug concentrations for each drug for individual donors. Body of the box plots represent first and third quartiles and red horizontal line represents the literature reported IC50. Error bars represent S.E.M. for each drug. Mechano-IC_50_ was calculated by fitting plot to a standard dose-response function: Signal=Bottom + (Top-Bottom)/(1+([drug]/IC50). g) Plot of MCATS signal (purple) and primary aggregation (PA, beige) signal obtained from donor platelets upon activating with ADP. Aggregometry data collected from a single donor. MCATS signal is the average from *n*=4 donors. The values were normalized to the signal obtained from the wells with 20 μM ADP.

We first tested MCATS as a function of the number of platelets seeded on the unzipping duplex surfaces. We found that MCATS detected tension signal from as few as 2000 platelets (**Figure S8**). We choose 2 × 10^6^ platelets in the following experiments because this concentration of platelets in a 96-well plate offered the strongest signal for the assay. Comparing MCATS signal on the unzipping and shearing duplex probes with 2 × 10^6^ human platelets showed that platelets produced more signal on the unzipping mode probes which is consistent with literature reports (**Figure S9**)Λ ^18^ In a second set of experiments, we incubated platelets with different drugs at room temperature for 30 minutes and plated 2 × 10^6^ treated human platelets in each well for 1hr and then performed MCATS.

For all the compounds tested, we observed a dose-dependent decrease in the 12 pN duplex rupture in both microscopy imaging of duplex ruptures as well as MCATS signal that was detected using a plate reader (**Figure 3c-f**, **Figure S10**). All drugs showed a significant drop in signal upon treating platelets. (*P*=0.0004, 0.009, <0.0001,0.005 for aspirin, Eptifibatide, 7E3 and Ticagrelor respectively) By fitting the plot of MCATS signal with a standard dose-response inhibition function (Signal=Bottom + (Top-Bottom)/(1+([drug]/IC_50_)), we determined the mechano-IC_50_ for aspirin, eptifibatide, 7E3 and Ticagrelor for each individual donor and plotted the data in **Figure 3c-3f**. MCATS was found to be robust as the signal generated from the same blood draw of the same donor was highly consistent (σ ~ 10%). However, we found donor-to-donor variability in mechano-IC50 which likely reflects the biological heterogeneity of the drug response especially in aspirin treated group. Importantly, the values we measured were consistent with literature precedent.^25–29^ Another parameter that could influence platelet forces is ADP agonist which is well known to trigger platelet activation and adhesion. We therefore tested the dose-response of platelets to agonist using MCATS and found increasing tension signal as a function of ADP. These results were further validated with light transmission aggregometry (LTA) which is commonly used to assess platelet function (**Figure 3g, Figure S11**). Aggregometry, however, requires 100-fold greater sample volumes and dedicated instruments which shows the advantage of MCATS in assessing platelet function with cellular forces. Other agonists such as TRAP and collagen were also tested and showed increasing tension signal as a function of dose of agonist (**Figure S12**). Because each measurement only requires 2 × 10^6^ platelets while 1ml of blood contains ~10^9^ platelets, a typical 5 ml blood draw can be used to run ~2,500 assays thus opening the door for massive screening to determine drug sensitivity in a personalized manner.

### MCATS detects platelet dysfunction and correlates with transfusion need in subjects following CPB

Finally, we investigated whether MCATS can be used to assess platelet dysfunction in cardiac patients using CPB. In a subset of patients (10-23%), CPB leads to severe postoperative bleeding requiring blood transfusion^30–32^. Past literature has demonstrated that CPB surgery and other extracorporeal circuits such as extracorporeal membrane oxygenation (ECMO) lead to changes in membrane glycoprotein GPIb-IX-V^33^ as well as integrin activation^34^ which results in platelet dysfunction.^35^ The likelihood of patient bleeding following CPB is typically assessed using platelet function assays such as impedance aggregometry and thromoboelastography (TEG). Despite their widespread use, such assays provide only a weak prediction of severe bleeding. For example, the PLATFORM clinical trial showed that impedance aggregometry has a 40% predictive value for severe bleeding^36^ while other studies showed that thromboelastography (TEG) has a 0.78 area under the curve (AUC) for predicting postoperative bleeding^37, 38^. This is likely because TEG and aggregometry are highly dependent on platelet count which can bias aggregation dynamics.^39^ The ability to anticipate the risk of developing coagulopathy due to platelet dysfunction more accurately is desirable as it could aid in determining the optimal timing of cardiac surgery for patients on antiplatelet agents, as well as targeting platelet transfusions to offer blood products to the patients who need it most. Given that bleeding risk is multifactorial, developing additional assays based on different transduction mechanisms may complement current techniques and offer improved capabilities to detect platelet dysfunction and bleeding risk.

We performed MCATS on *n*=6 healthy subjects as well as *n*=7 cardiac patients pre- and post-CPB (**Figure 4a**). The demographic information and related health information for these subjects can be found in **Table S3**. All CPB patients were tested using TEG (using a TEG6S system) and aggregometry in the clinic pre- and post- surgery, and these measurements were benchmarked against MCATS. Note that due to low count platelet, one subject did not have an accompanying aggregometry measurement. The MCATS score for each donor, was averaged from *n*=3 or 4 replicate measurements where 2×10^6^ platelets were seeded in a well and the fluorescence intensity was measured at *t* = 1 hr after seeding. In general, healthy donors (*n*=6) showed similar MCATS tension signal which was greater than that for CPB donors prior to surgery. This is likely because many of the CPB patients had underlying health conditions, but this premise is yet to be validated. After CPB, subjects showed a significantly reduced MCATS signal (decrease of 35% ±18%, *P*=0.0002) that ranged from 48% to 17% (**Figure 4b**). Within 24 hrs after CPB, 5 out of the 7 patients required platelet transfusion to minimize bleeding. We hypothesized that the magnitude of MCATS signal reduction (mechanical dysfunction) is an indicator of the severity of platelet dysfunction, and hence we compared the change in MCATS signal (%) to the number of units of transfused platelets (**Figure 4c**). We found that the number of transfused platelets given to the patient within 24 hrs after surgery were strongly positively correlated with the drop in platelet MCATS signal (**Figure 4c, Pearson’s r= 0.83, *P*=0.020**). As expected, aggregometry analysis (with 10μM of ADP) of the same samples showed a decrease of 47% ± 18%, *P*=0.05 in primary aggregation following CPB. TEG showed a decrease of 29% ± 14%, *P*=0.001 in maximum aggregation (MA) for the same cohort (**Figure 4d, 4f**). Note that one of the CPB subjects showed an increase in aggregometry value following surgery which further underscores the variability in aggregometry. When the aggregometry and TEG signal change was compared to the number of platelet units transfused, we found a weakly positive correlation for aggregometry and a strong correlation for TEG (**Figure 4e, 4g, Pearson’s r= 0.16, *P*=0.32 for aggregometry and Pearson’s r= 0.86, *P*=0.012 for TEG**). Our pilot study strongly supports the premise that the mechanical forces generated by platelets offer a robust metric to quantify platelet function that is comparable to the methods being used in the operating room such as TEG and thus may offer a complementary indicator of post-op bleeding risk following CBP.

**Figure 4.**
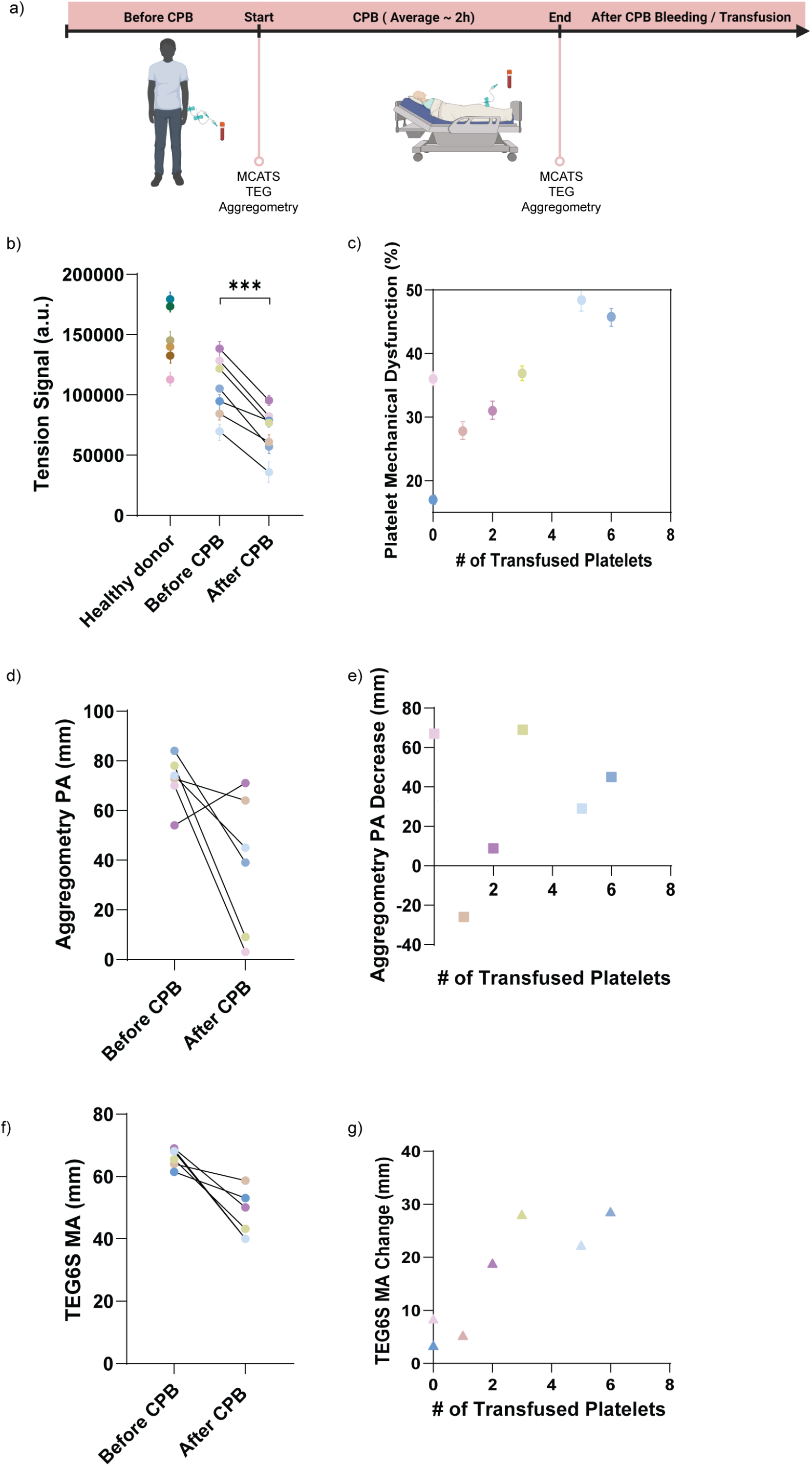
MCATS detects platelet dysfunction for patients following CPB. a) Schematic of workflow using MCATS to assess platelets dysfunction following CPB. b) Plot of MCATS platelet signal for healthy donors (n = 6), pre-CPB and post-CPB subjects (n = 7). Error bars represent S.D. from n= 3 or 4 measurements for the same platelets. Significance is calculated with two tailed student t-test with *P*=0.002. c) Plot of MCAT reduction in signal (%) against the number of platelets transfusions given to the subject within 24 hrs of CPB. Data points are color coded for each subject. Error bars represent S.D. of n= 3 or 4 replicates for each sample. Pearson’s correlation coefficient r= 0.83 with *P*=0.020. d) Plot of aggregometry PA (mm) for subjects pre-CPB and post-CPB. e) Plot of aggregometry PA decrease (mm) vs platelet transfusion needs within 24 hrs of CPB. Note that one patient lacked aggregometry data because of severe hemolysis. Pearson’s correlation coefficient r= 0.16 with *P*=0.32. f) Plot of TEG MA (mm) for subjects pre-CPB and post-CPB. g) Plot of TEG MA decrease (mm) vs platelet transfusion need. Pearson’s correlation coefficient r= 0.86 with *P*=0.012.

We also performed controls to validate MCATS measurement for subjects following CPB. Following standard of care procedures, patients were treated with heparin prior to surgery to minimize blood clotting and this was neutralized with protamine post operation. To confirm that the decrease in MCATS signal is not due to inhibition of heparin, we performed MCATS on controlled platelets that were treated with heparin and then neutralized with protamine. The results showed that heparin led to a slight decrease in MCATS signal, but this inhibition was fully reversed upon protamine treatment (**Figure S13**). We also tested the mechano-IC50 of aspirin and Ticagrelor for the CPB donors before and after surgery. The results showed that the surgery did not influence the mechano-IC_50_ significantly (**Figure S14**), which confirms that MCAT is insensitive to platelet count.

## Discussion

The current state of the art tools for measuring platelet function are based on aggregometry and viscoelastic assays (TEG)^40^ which are routinely used in clinical practice. Aggregometry measures the light transmission of platelet rich plasma sample(~1mL) using a dedicated instrument that applies shear as well a specific agonist such as ADP and TRAP. Aggregometry records multiple parameters including the kinetics of aggregation to infer clotting functions. Similarly, TEG uses a dedicated instrument that records the drag forces in whole blood samples which measures the kinetics of clot formation and viscoelastic properties of clots. The limitations of these assays are multifold. First, unlike MCATS which requires a standard well plate reader that is already used for most ELISA assays, TEG and aggregometry assays require dedicated instruments which limits widespread adoption. Secondly, LTA and TEG require ~mL quantities of blood for each measurement which may not seem significant, but this prohibits running triplicates and performing drug screens. Even more importantly, TEG and LTA assays have limited sensitivity given that these assays require platelet aggregation which is highly dependent on platelet count, coagulation factors such as fibrinogen level and antiplatelets agents usage.^41–43^ Patients with higher platelet counts will tend to form more stable aggregates regardless of the functional activity of each individual platelet^44^ and patient with a low platelets count due to surgery or underlying conditions have difficulty to run TEG or aggregometry assays. According to the PLATFORM study, aggregometry shows limited predictive value in identifying post operative bleeding following CPB.^36^ Indeed, in our work, we encountered one patient with a low platelet count which would present a significant challenge for TEG but this did not impact the MCATS assay.

One potential clinical use of MCATS may be in predicting severe bleeding. The universal definition for perioperative bleeding in adult cardiac surgery stratifies patients into five classes, where class 1 and 2 are defined as insignificant, and mild bleeding, while class 3, 4, and 5 indicate moderate, severe, and massive bleeding, respectively.^45^ The definition is based on a number of factors including blood loss and allogeneic blood product transfusion. We followed this definition and classified CPB patients based on clinical observations. We found that patients with moderate, severe, or massive bleeding (class 3, 4, 5) showed a more significant drop in MCATS score compared to patients with mild or insignificant bleeding (**Figure S15)**. Because MCATS exclusively quantifies platelet function, more studies are needed for understanding the correlation between platelet function and bleeding risk.

A unique merit of MCATS is that it measures molecular forces directly using conventional plate reader and it uses Cas12a amplification to boost the signal for fast and sensitive readout. As a result, MCATS only requires ~5 μL of blood or less, and hence a typical blood draw (~5 mL) is enough to run 1000 assays. Antiplatelet therapies are commonly prescribed to prevent acute arterial thrombosis. However, antiplatelet therapy assessment based on aggregometry has not shown improved outcomes and better benefit/risk for cardiovascular patient.^46^ MCATS can generate personalized dose-response curves for specific drugs rapidly to optimize treatment in a dynamic manner, which may increase the sensitivity and specificity required to guide personalized platelet therapy. So far, we measured the forces applied by GPIIb/IIIa which mediates platelet adhesion and aggregation to assess platelet functions. Other platelet forces mediated by GPIb-IX-V and GPVI can potentially be investigated by attaching different ligands/proteins on DNA tension probe to understand the platelet forces with different receptors under shear flow or in different hemostasis states.^47^ It is also worth noting that live and active cells are required to perform MACTS, and we confirmed that lyophilized platelets (fixed platelets with intact protein structure) failed to generate MCATS signal (**Figure S16**).

In MCATS experiments, different platelet handling procedures may alter platelet response. Therefore, we tested different platelet purification procedures and the results indicated that MCATS can detect tension signal from purified platelets, platelet rich plasma (PRP) and whole blood sample. However, whole blood showed decreased tension signal in MCATS because the high abundance of red blood cells blocked access to the chip surface. We anticipate that applying mild flow conditions would better allow performing MCATS in whole blood. PRP showed slightly decreased MCATS signal compared to that when using purified platelets. Advantages of using PRP include simplifying blood processing steps and decreasing preparation time to run the assay (**Figure S7**). Other advantages of using PRP include maintaining the physiological drug dose found in the blood which is important for assessing drugs that have short half-life or rapid off-rates. Conversely, the use of purified platelets for MCATS enhanced the signal and provided a more direct evaluation of platelet function without interference from soluble factors.

Another important parameter to consider in MCATS is the assay time. We monitored MCATS signal in platelet and fibroblast experiments with different amplification times. The results showed that 30min amplification provides sufficient signal, but signal could be further enhanced at 60 min. Further increasing amplification time did not lead to significantly improved signal to noise ratio (**Figure S17**). These results indicate that MCATS has potential as a point-of-care assay given the 30 min or 60 min response time.

In summary, we developed an ultrasensitive fluorescence-based assay to rapidly measure cell receptor mediated tension. When cell receptors apply sufficient force, the DNA duplex tension sensor denatures exposing single stranded DNA. We amplify the peeled DNA using Cas12a to produce ~10^4^ fluorophores in response to each 12 pN event, which allows high-throughput detection of cellular tension without requiring any dedicated hardware. Instead, the assay uses modified 96 well plates that are read out in a conventional plate reader found in all clinical chemistry laboratories. MCATS offers an important step toward taking assays that are used in molecular biophysics and mechanobiology toward translational applications.

## Methods

### Materials

Cy3B-NHS ester (PA63101) was purchased from GE Healthcare Life Sciences (Pittsburgh, PA). Atto647N-NHS ester (18373) was purchased from Sigma Aldrich (St. Louis, MO). Cyclo[Arg-Gly-Asp-d-Phe-Lys(PEG-PEG)] (PCI-3696-PI) (cRGD) was acquired from Peptides International (Louisville, KY). Streptavidin (S000-01) was purchased from Rockland-Inc (Pottstown, PA). μ-Slide VI0.4 6-channel slides (80606) and 25 mm x 75 mm glass coverslips (10812) were purchased from Ibidi (Verona, WI). ProPlate® Microtiter (204969) were purchased from Thermo-Fisher Scientific. 200 proof ethanol (100%, #04-355-223) was purchased from Fischer Scientific. (Waltham, MA) N-hydroxyl succinimide-5 kDa PEG-biotin (NHS-PEG-biotin, HE041024-5K) was purchased from Biochempeg (Watertown, MA). 3-Aminopropyl triethoxysilane (APTES, 440140, 99% purity) and adenosine 5’-diphosphate (ADP, A2754, 95% purity), and KKO (Lot: XE3598104) were purchased from Sigma-Aldrich. Ticagrelor was acquired from Selleck Chemistry (Houston, TX). 7E3 (Abciximab, Lot: GR3422387-2) was purchased from Abcam. Apyrase and LbCas12 were purchased from New England Biolabs (Ipswich, MA). All DNA oligonucleotides used in this work are listed in **Table S1** and were custom synthesized by Integrated DNA Technologies (Coralville, IA). crRNA was custom synthesized by Dharmacon Inc.(Lafayette, CO). All other reagents and materials (unless otherwise stated) were purchased from Sigma-Aldrich and used without purification. All buffers were prepared with 18.2 MΩ nanopure water.

### Instruments

We used a Nikon Eclipse Ti microscope, operated by Nikon Elements software, and equipped with a 1.49 numerical aperture (NA) CFI Apo ×100 objective, perfect focus system, a TIRF laser launch, a Chroma quad cube (ET-405/488/561/640 nm Laser Quad Band) and an RICM (Nikon: 97270) cube for this work. Bulk fluorescence measurements were conducted using a Synergy H1 plate reader (Bio-Tek) using fluorescence filter sets. All ultrapure water was obtained from a Barnstead Nanopure water purifying system (Thermo Fisher) that indicated a resistivity of 18.2 MΩ. Nucleic acid purification was performed using a high-performance liquid chromatography (HPLC, Agilent 1100) equipped with a diode array detector. Microvolume absorbance measurements were obtained using a Nanodrop 2000 UV-Vis Spectrophotometer (Thermo Scientific). Vacufuge Plus (Eppendorf) was used to remove solvent from HPLC purified sample. Electro-spray ionization (ESI) mass spectrometry identification of product was performed with an Exactive™ Plus Orbitrap Mass Spectrometer. MJ Research PTC-200 Thermal Cycler was used to anneal and hybridize DNA. Platelet counts were determined on a Poch-100i hematology analyzer.

### Surface Preparation

MCATS surface preparation was adapted from previously published protocols.^18, 48^ Briefly, rectangular glass coverslips (25 × 75 mm, Ibidi) were rinsed with water and sonicated for 20 minutes in DI water and then this was repeated for another 20 minutes in ethanol. The glass converslips were then cleaned with piranha solution which was prepared using a 1:3 mixture of 30% H_2_O_2_ and H_2_SO_4_.^49^ WARNING: Piranha solution becomes very hot upon mixing, and is highly oxidizing and may explode upon contact with organic solvents. Please handle with care. Slides were then washed 6 times in ultrapure water, followed by 4 successive washes using ethanol. In a separate beaker of ethanol, slides were reacted with 3% v/v APTES at room temperature for 1 h. Coverslips were then washed 6 times with ethanol, baked in an oven for 20 minutes at 80 °C. Subsequently, the amine terminal groups were coupled to NHS-PEG-biotin (3% w/v) by placing 200 uL of a freshly prepared 6 mM NHS-PEG-biotin solution in ultrapure water between two slides for 1 hr. Next, slides were washed 3 times with ultrapure water, dried under N_2_ gas, and then stored at −30°C for up to 2 weeks or longer before use. At the day of imaging, the 5kDa PEG-biotin substrate was adhered to a ProPlate microtiter 96-well plate housing with an adhesive bottom. Wells were then incubated with 50 μg/ml (830 nM) streptavidin in 1 × PBS for 1 hr. Wells were then washed with 1XPBS and incubated with 100 nM DNA probe solutions for 1 hr. Finally, the wells were washed with 1x PBS.

### DNA Hybridization and gRNA/Cas12a binding

DNA oligonucleotides were hybridized at 100 nM in a 0.2 mL PCR tube before incubating on the surface. Using a MJ Research PTC-200 thermocycler, DNA was first heated to 90°C and then cooled at a rate of 1.3°C per min to 35°C. gRNA and Cas12a were incubated for 10min at 37 °C at 500nM in a 0.2 mL PCR tube just before adding to the surface and stored on ice for maximum preservation of activity.

### Oligo dye/ligand coupling and purification

All sequences of DNA strands used in this work are provided in **Table S1**. To generate the dye-labeled bottom strand, 10 nmoles of amine-modified DNA was reacted overnight at 4°C with a 20_x_ excess of Cy3B-NHS or ATTO 647N-NHS dissolved in 10 μL DMSO. The total volume of the reaction was 100 μL and this was composed of 1x PBS supplemented with 0.1M NaHCO_3_. Then a P2 size exclusion gel was used to remove unreacted dye. The product **1** or **2** (**Fig. S5**) was then purified by reverse phase HPLC using an Agilent Advanced oligo column (Solvent A: 0.1M TEAA, Solvent B: acetonitrile; starting condition: 90% A + 10 % B, 1%/min gradient B, Flow rate: 0.5 mL/min) (**Fig. S5**).

To generate the cRGD-modified top strand (product **3** in **Fig. S5**), 100 nmoles of c(RGDfK (PEG-PEG)) was reacted with ~ 200 nmoles of NHS-azide in 15 uL of DMSO overnight at 4°C. Product **3** was then purified via reverse phase HPLC using a Grace Alltech C18 column (solvent A: water + 0.05% TFA, solvent B: acetonitrile + 0.05% TFA; starting condition: 90% A + 10 % B, 1%/min; flow rate: 1 mL/min).

Purified product **3** was ligated to the BHQ2 top strand via 1,3-dipolar cycloaddition click reaction. Briefly, 5 nmoles of alkyne-modified top strand was reacted with ~70 nanomoles of product **3**. The total reaction volume was 50 μL, composed of 0.1 M sodium ascorbate and 0.1 mM Cu-THPTA for 2h at RT. The product **4** was then purified with a P2 size exclusion column, and then purified with reverse phase HPLC using an Agilent Advanced oligo column (solvent A: 0.1M TEAA, solvent B: acetonitrile; starting condition: 90% A + 10 % B, 0.5%/min gradient B, flow rate: 0.5 mL/min) (**Fig. S5**).

Concentrations of purified oligonucleotide conjugates were determined by measuring their absorption at λ=260 nm on a Nanodrop 2000 UV-Vis Spectrophotometer (Thermo Scientific). ESI- mass spectrometry was performed to validate all oligonucleotide products, and the results are listed in **Table S2**.

### Solution based Cas12a amplification and plate reader readout

Cas12a reactions reported in **Figure 1e** were performed at 37°C for 1h in 1 × PBS supplemented with 10 mM MgCl2. For these measurements, we maintained a total DNA (60 T bottom strand, **Table S1**) solution concentration of 100 nM but tuned the ratio between the blocked DNA and unblocked activator. We prepared freshly primed Cas12a by mixing with gRNA at 1:1 ratio. The primed Cas12a-gRNA complex was then mixed at 20 nM concentration with 100 nM fluorogenic reporter to the well. Then we measured the final fluorescence intensity using a plate reader using a filter set (Ex/Em = 540/590 nm for reporter channel) after allowing the reaction to proceed for 1h.

### Human platelet handling and ethics agreement

Blood was collected at Emory Hospital from consented patients/volunteers. For patients, blood was collected from arterial line into 5ml 3.2% citrate tubes (9:1 v/v) at baseline and after CPB (after protamine treatment). For healthy volunteers, blood was collected by venipuncture into citrate tubes as above. Complete blood count (CBC) was performed using Poch-100i hematology analyzer (Sysmex Corp, Kobe, Japan) to evaluate platelet count (PLT). The sample was then centrifuged for 12 min at 140 g (with 0.02U Apyrase). Then PRP was separated and centrifuged for 5 min at 700 g with 3 μM PGE-1. The platelets were then resuspended in Tyrodes buffer with 3μM PGE-1 and centrifuged for 5 min at 700 g. Finally, platelets were resuspended in Tyrodes buffer. It is worth noting that apyrase is important for the platelet purification procedure to prevent hemolysis-triggered platelet aggregation.

Ethics: This project has approval from the Emory University Ethics Committee to obtain samples from participants and these ethical regulations cover the work in this study. Written informed consent was obtained from all participants.

### Cell culture

NIH/3T3 fibroblasts were cultured according to ATCC guidelines. Briefly, cells were cultured in DMEM supplemented with 10% bovine calf serum (v/v, purchased from Gibco) and penicillin/streptomycin. Cells were passaged every 2-3 days as required.

### Mechano-Cas12a assisted tension sensor

MCATS was performed on activated substrates as described in the surface preparation section. First, cRGDfK-labelled concealed activator probes were incubated on biotin surfaces in 1x PBS buffer for 1h. The wells were washed with 1x PBS. Then, cells were added and allowed to spread on the surfaces for 1h in specific media (DMEM supplemented with 1% serum for NIH/3T3 cells and Tyrodes buffer with 10 μM ADP for platelets). Surfaces were imaged at approximately 1hr after seeding using epifluorescence and TIRF microscopy to visualize and quantify mechanically exposed activators. Wells were then supplemented with 10 mM MgCl2 and subsequently, 20 nM gRNA/Cas12a complex and 100 nM reporter DNA were mixed and added to the well to initiate the Cas12a amplification reaction with mechanically exposed activator. After 1h of triggering the Cas12a reaction, fluorescence intensities of wells were measured with a Bio-Tek® Synergy H1 plate reader (Ex/Em = 540/590 nm for reporter channel). Note that we always included positive and negative control wells to help benchmark the MCATS signal. The positive control wells were comprised of 100% surface tethered activator oligonucleotides while the negative control was the blocked activator.

### Dose-dependent inhibition of receptor mediated tension

For dose-dependent inhibition of experiments, the cell density of 3T3 fibroblast was first characterized with a hemocytometer. 25×10^3^ cells were incubated with different concentrations of inhibitor in the cell culture incubator for 30 min before plating onto 96 well plates. Afterwards, cells were incubated for 1h to promote cell adhesion. Then the MCATS protocol was followed to achieve amplification and quantification.

For platelet MCATS measurements, human platelets were purified and stored at room temperature for at least 30 min before running experiments. For the drug treatment measurements, platelets were treated with each drug for 30 min before seeding onto 96 well-plates. 10 μM ADP was added to the well to promote cell activation and adhesion. Then platelets were incubated at room temperature for 1h. The same MCATS protocols that was used for fibroblasts was also followed to achieve amplification and quantification of force generation by platelets.

### Microscopy imaging

For MCATS experiments, Images were acquired on a Nikon Eclipse Ti microscope, operated by Nikon Elements software, a 1.49 NA CFI Apo 100x objective, perfect focus system, and a total internal reflection fluorescence (TIRF) laser launch with 488 nm (10 mW), 561 nm (50 mW), and 638 nm (20 mW). A reflection interference contrast microscopy (RICM) (Nikon: 97270) cube and a Chroma quad cube (ET-405/488/561/640 nm Laser Quad Band) were used for imaging. Imaging was performed on 96 well plates and glass coverslips using DMEM as cell imaging media for 3T3 cells and Tyrode’s buffer for platelets. All imaging data was acquired at room temperature.

### Thromboelastography

TEG measurements were obtained in the operating room using the TEG 6S system (Haemonetics, Braintree, MA) according to manufacturer instructions. This is a fully-automated point of care system shown to have excellent agreement with the TEG 5000 (Haemonetics, Braintree, MA), which has been widely used in cardiac surgery for purposes of guiding hemostatic blood product transfusions.^50^ Briefly, 2.7 ml of whole blood was collected from the subject’s preexisting arterial line into 3.2% citrated tubes. Following the recommended wait time of 10 minutes, approximately 400 μl of whole blood was pipetted into the sample port of a global hemostasis cartridge (Haemonetics, Braintree, MA). The parameter of interest was the maximum amplitude (MA) of the standard kaolin activated channel. The MA reflects the strength of platelet-fibrinogen interactions.

### Light Transmission Platelet Aggregometry

To isolate PRP, blood collected from volunteers/patients was centrifuged at room temperature at 100 × g for 12 min in Megafuge 16R (ThermoScientific, Waltham, MA) equipped with swinging bucket rotor. The PRP layer was removed and PLT count was obtained and if necessary adjusted to less than 400,000/μl. Next, samples were re-centrifuged at 2000xg for 20 min to obtain platelet poor plasma (PPP). LTA was accomplished using PAP-8E profiler (Biodata Corp, Horsham, PA) prewarmed to 37°C. The blank (0% aggregation) was set with PPP. For the testing, ADP and TRAP-6 were used as platelet activators. The final concentrations used were 20, 10 μM for ADP and 10 μM for TRAP. All LTA testing was performed according to manufacturer’s directions. % Aggregation (PA), slope (PS), area under the curve (AUC), lag phase (LG), disaggregation (DA), maximum aggregation (MA) and final aggregation (FA) were recorded.

### Statistics and reproducibility

P-values were determined by two tailed student’s t-test using GraphPad Prism8. Mechano-IC_50_ is calculated by fitting plot to a standard dose-response function: Signal=Bottom + (Top-Bottom)/(1+([drug]/IC50) using GraphPad Prism8. Each MCATS readout is typically replicated with 2 - 4 wells on the same plate for the same condition. For CPB patient tension signal, all data was acquired along with positive and negative control wells to help benchmark signal. The positive control wells were comprised of 100% surface tethered activator oligonucleotides while the negative control was the blocked activator without platelet seeding. MCATS signal for each run was normalized to the positive control wells also conducted in each experiment and then multiplied by the average intensity of the positive wells collected from past positive control runs. This normalization process helped to minimize the influence of variance in enzyme activity (batch to batch variability) along with variability in the surfaces and other sources of experimental noise.

### Reporting summary

Further information on research design is available in the Nature Research Reporting Summary linked to this article.

### Data availability

The data supporting the results in this study are available within the paper and its Supplementary Information. Source data are provided with this paper. All raw and analyzed datasets generated during the study are available from the corresponding author on request. De-identified patient data are available from the corresponding author, subject to IRB approval.

## Acknowledgements

We acknowledge support from NIH 5R01GM131099-04, NSF DMR 1905947. We thank Dr. Hiroaki Ogasawara for helping run ESI-MS to validate oligonucleotides. We thank Dr. Laura Downey and Dr. Cheryl Maier for helpful discussions.

## Author Contributions

Y.D. and K.S. conceived the project. Y.D. designed experiments, analyzed data, and compiled the figures. F.S. helped with TEG, aggregometry experiments and related discussions. Y.H., W.C., R.L. M.M. helped with platelets purification and related discussion. Y.K. helped design experiments. F.S. and R.S. helped designed clinical studies and obtained related samples. Y.D. and K.S. wrote the manuscript. All authors helped revise the manuscript.

