## Supplementary Information for "Mechano-Cas12a Assisted Tension Sensor (MCATS) for Massively Amplified Cell Traction Force Measurements"

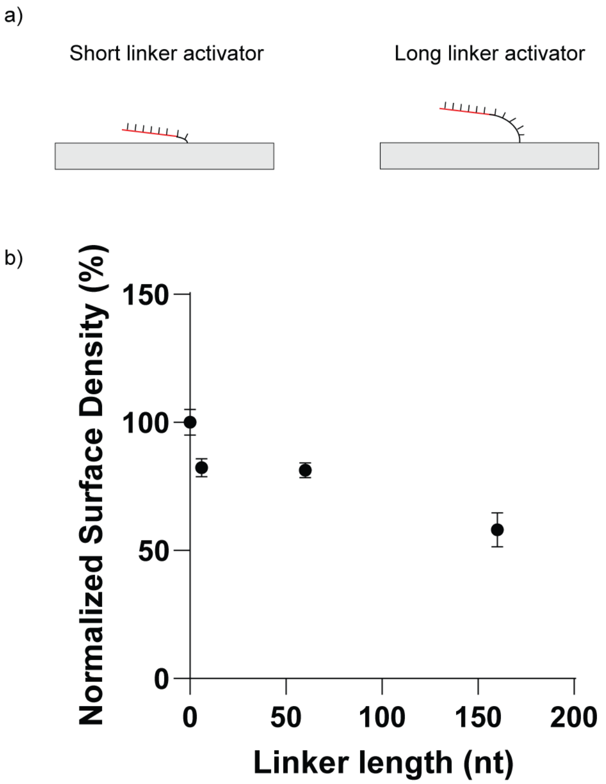

**Figure S1. Comparison of surface density with different length of linkers.** A) Scheme showing activators with different linker length on the surface. B) Plots of normalized surface density vs linker length. All activators are incubated on surface for 1hr at 100nM. Surface density is normalized to activator without linker. Error bar represents S.E.M from five independent surfaces.

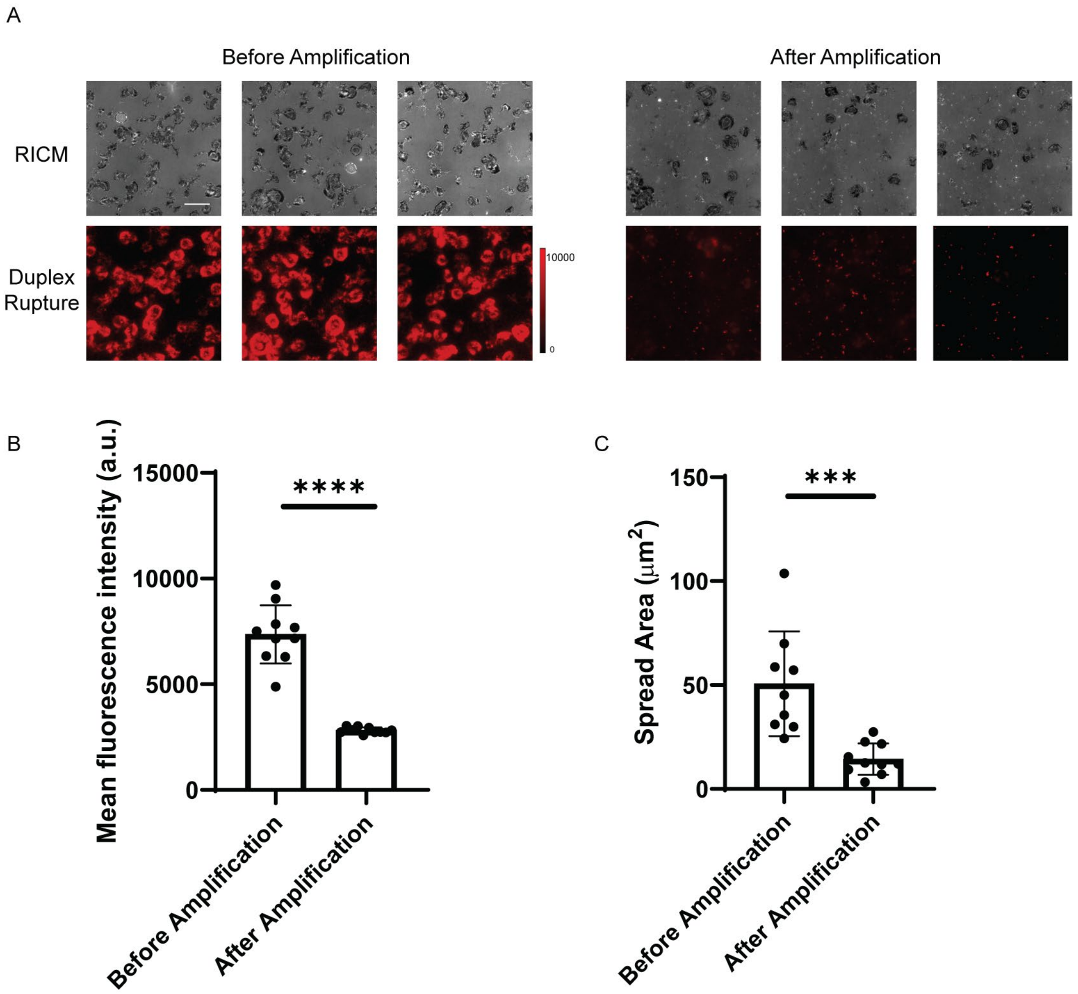

**Figure S2. Cas12a auto-cleavage of surface-tethered activator after activation.** A) Representative RICM, and duplex rupture (red) fluorescence images after platelets were incubated on concealed activator surface. The images compare the RICM cell spread area and duplex rupture signals before and after Cas12a/gRNA and reporter DNA were added for 1hr. Scale bar = 12 μm. B) Plot of mean fluorescence beneath platelets individual before and after amplification. Significance is calculated with two tailed student t test, $P<0.0001$. Value is not background subtracted. Error bar represents S.D from 10 individual platelets C) Plot of spread area beneath individual platelets determined from RICM before and after amplification. Significance is calculated with student t test, $P=0.0004$. Error bar represents S.D from 10 individual platelets

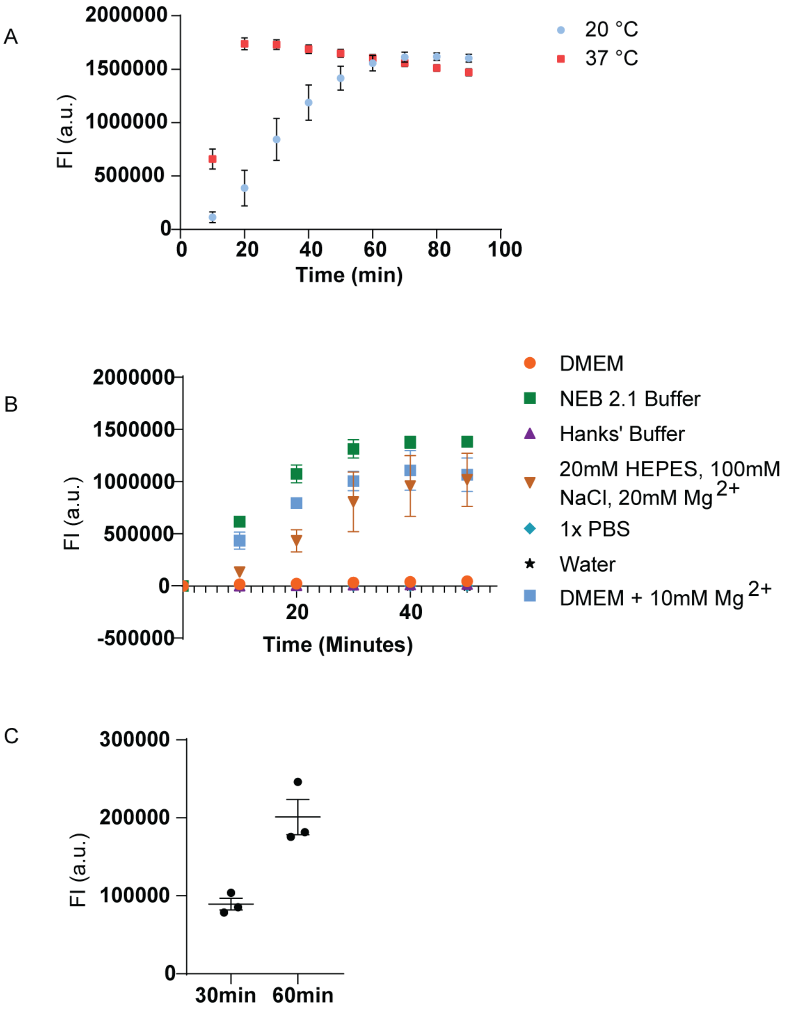

**Figure S3. Optimization of MCATS parameters of temperature, buffer, and reaction time.** A) Plot showing time-dependent fluorescence for Cas12a mediated hydrolysis of reporter DNA at 20 and 37 °C. The reaction was run with 100 nM soluble activator, 20 nM gRNA/Cas12a complex, and 100 nM reporter DNA. Results show that Cas12a activity is greater at 37 °C. Error bar represents S.E.M. obtained from three independent experiments. B) Plots of Cas12a activity in different buffers. We found that Cas12a is markedly less active in standard cell culture media compared to that of NEB buffer 2.1 which is likely due to decreased Mg ion concentration in cell media. Spiking cell culture media with 10 mM Mg2+ rescued Cas12a activity. Error bar represents S.E.M. obtained from three independent experiments. C) Plots of fluorescence signal at 30 and 60 min after triggering MCATS with Cas12a and reporter determined from human platelets (2*10^6). Error bar represents S.E.M. obtained from three independent experiments. Results indicate that 1hr amplification provides improved signal in cell experiments.

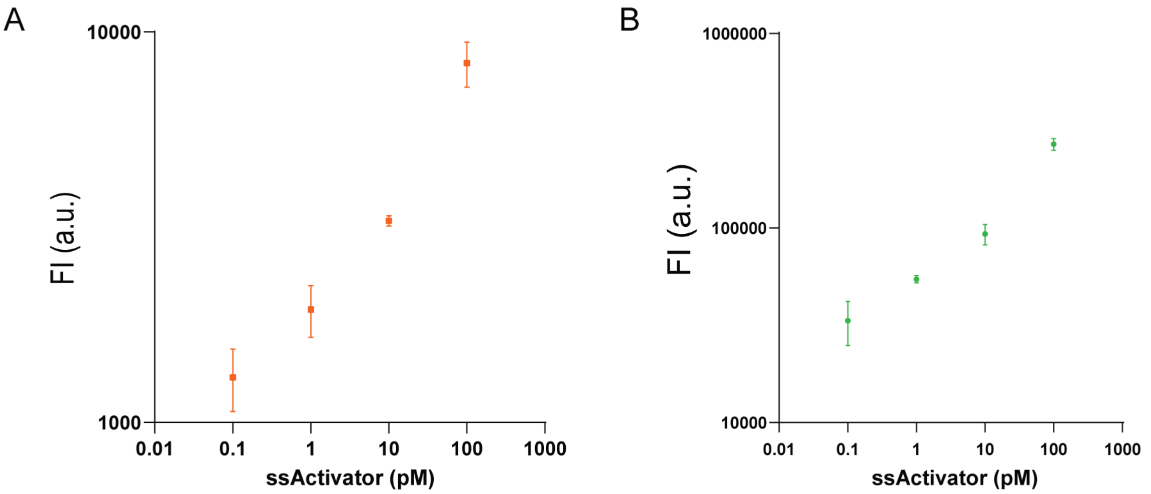

**Figure S4. Comparing MCATS with two fluorogenic ssDNA substrates.** A) 20 nM Cas12a-gRNA complex and 100 nM reporter DNA (/5ATTO565N/TT ATT /3BHQ_2/) of was added to various concentration of ssDNA and mixed with concealed activator for a total concentration of 100nM in solution, plot of fluorescence intensity-ssActivator concentration. Error bar represents S.E.M. obtained from three independent experiments. S/N (signal to noise ratio) = 35 for 100pM ssActivator. Noise was calculated from standard deviation of the blank B) 20 nM Cas12a-gRNA complex and 100 nM reporter DNA (/56-FAM/TT ATT /3IABkFQ/) of was added to 100 nM ssDNA and concealed activator in solution, plot of fluorescence intensity-ssActivator concentration. S/N = 18. The better S/N of Atto 565N reporter is chosen for more sensitive detection in later experiments.

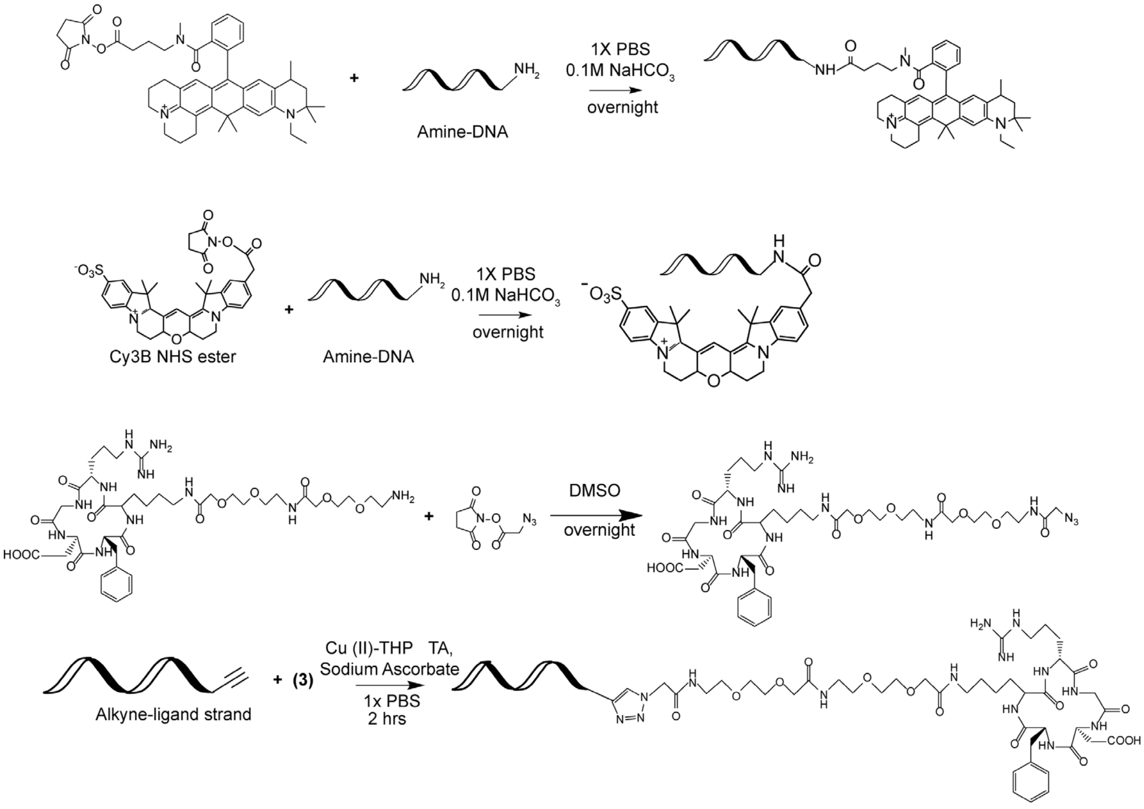

(1)

(2)

(3)

(4)

**Figure S5: Modified oligonucleotides.** Chemical structures and reactions of oligonucleotides, dye NHS esters and cRGDfk peptides.

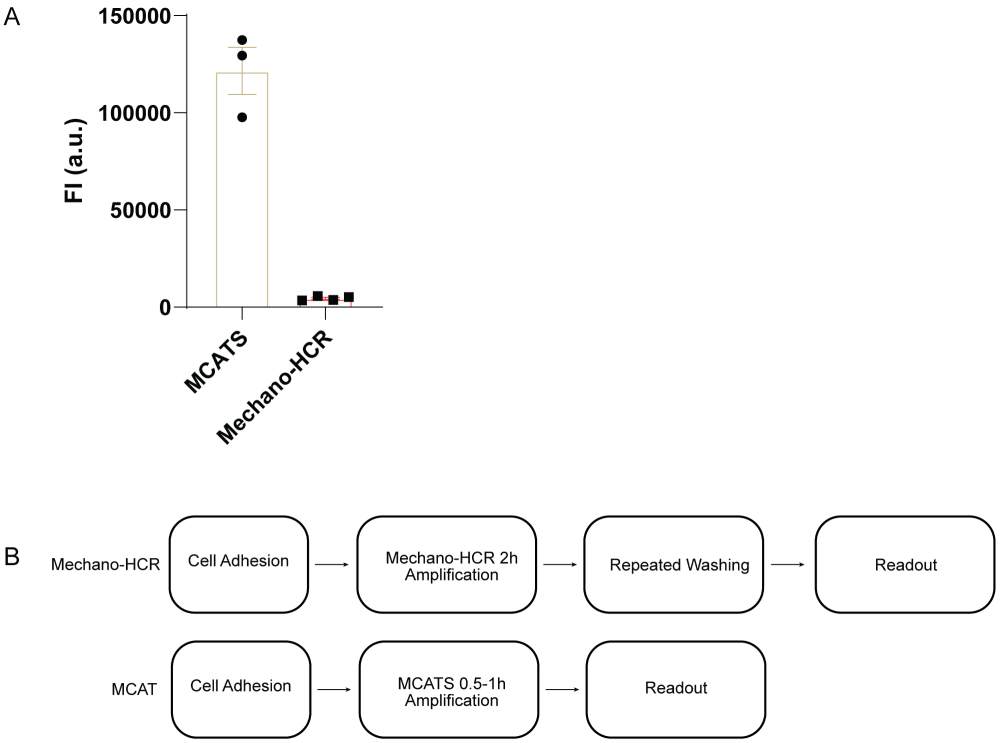

**Figure S6: Comparison between MCATS and Mechano-HCR.** A) Comparison of fluorescent signal of MCATS and Mechano-HCR amplified tension signal in the same day experiment with same number of cells (20000 NIH/3T3) seeded on the same surface and measured from same plate reader. B) Comparison of workflow of MCATS and Mechano-HCR. MCATS requires less steps in sample handling and less time in incubation

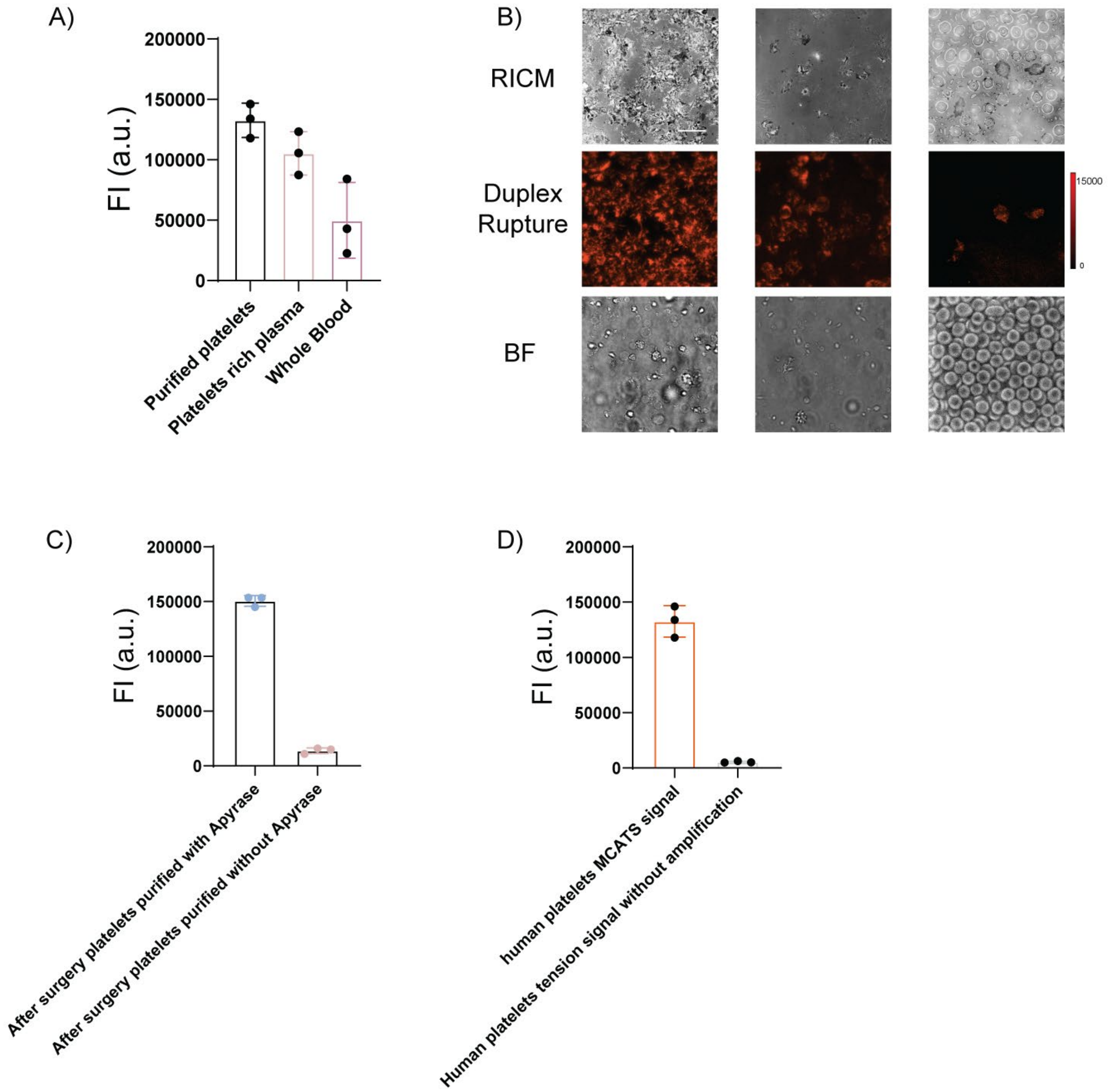

**Figure S7: Platelets handling optimization.** A) Comparison of MCATS signal using purified platelets, platelets rich plasma and whole blood. Error bar represents S.D from three wells with same patient sample. B) Representative of RICM, duplex rupture (red) fluorescence images and bright field images of tension signal when seeding different samples on concealed activator surface for 1hr. Scale bar = 12 μm. C) Comparison of MCATS signal in one experiment with platelet purification with/without Apyrase. In some post-surgery samples, patients' blood showed hemolysis during centrifugation, which required Apyrase to prevent platelets aggregation during purification. Error bar represents S.D from three wells with same patient sample. D) Comparison of MCATS and non-amplified duplex rupture signal using plate reader, indicating MCATS amplification is crucial for detecting platelets tension signal. Error bar represents S.E.M. from three independent experiments.

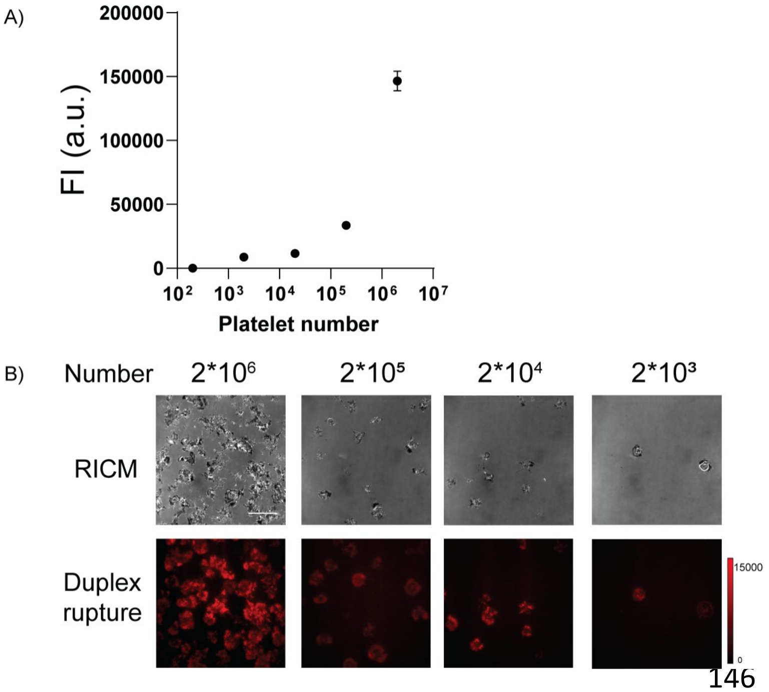

**Figure S8: Measuring MCATS signal with different number of platelets seeded on surface.** A) Comparison of MCATS signal using purified platelets, platelet rich plasma, and whole blood. Error bar represents S.E.M. from three independent experiments. B) Representative of RICM, duplex rupture (red) fluorescence images and bright field images of tension signal when seeding different number of cells on concealed activator surface. Scale bar = 10 μm

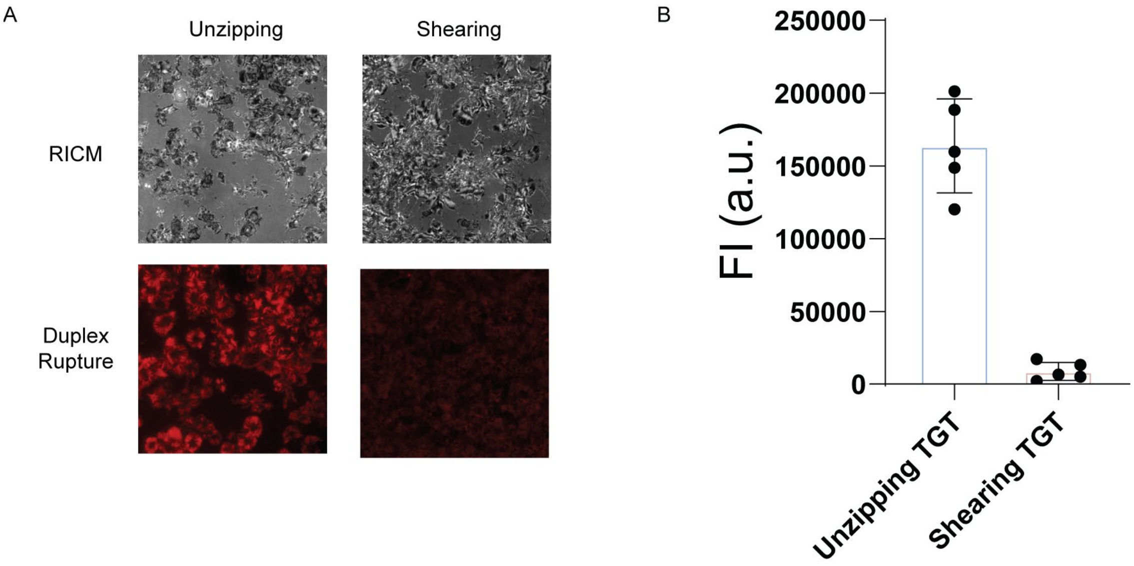

**Figure S9: Measuring human platelets tension with shearing and unzipping DNA tension probes.** A) Representative RICM and fluorescence images of cells cultured on $T_{tol}$= 12 pN and $T_{tol}$= 56 pN surfaces for 1hr. Scale bar = 12 μm. The red color is emission from Atto647N tagging the activator. The intensity bar next to each fluorescence image shows the absolute signal intensity for each image. B) Plots showing MCATS signal of same number of cells seeded on $T_{tol}$= 12 pN and $T_{tol}$= 56 pN surfaces. Error bars represent S.E.M. from n=3 independent experiments.

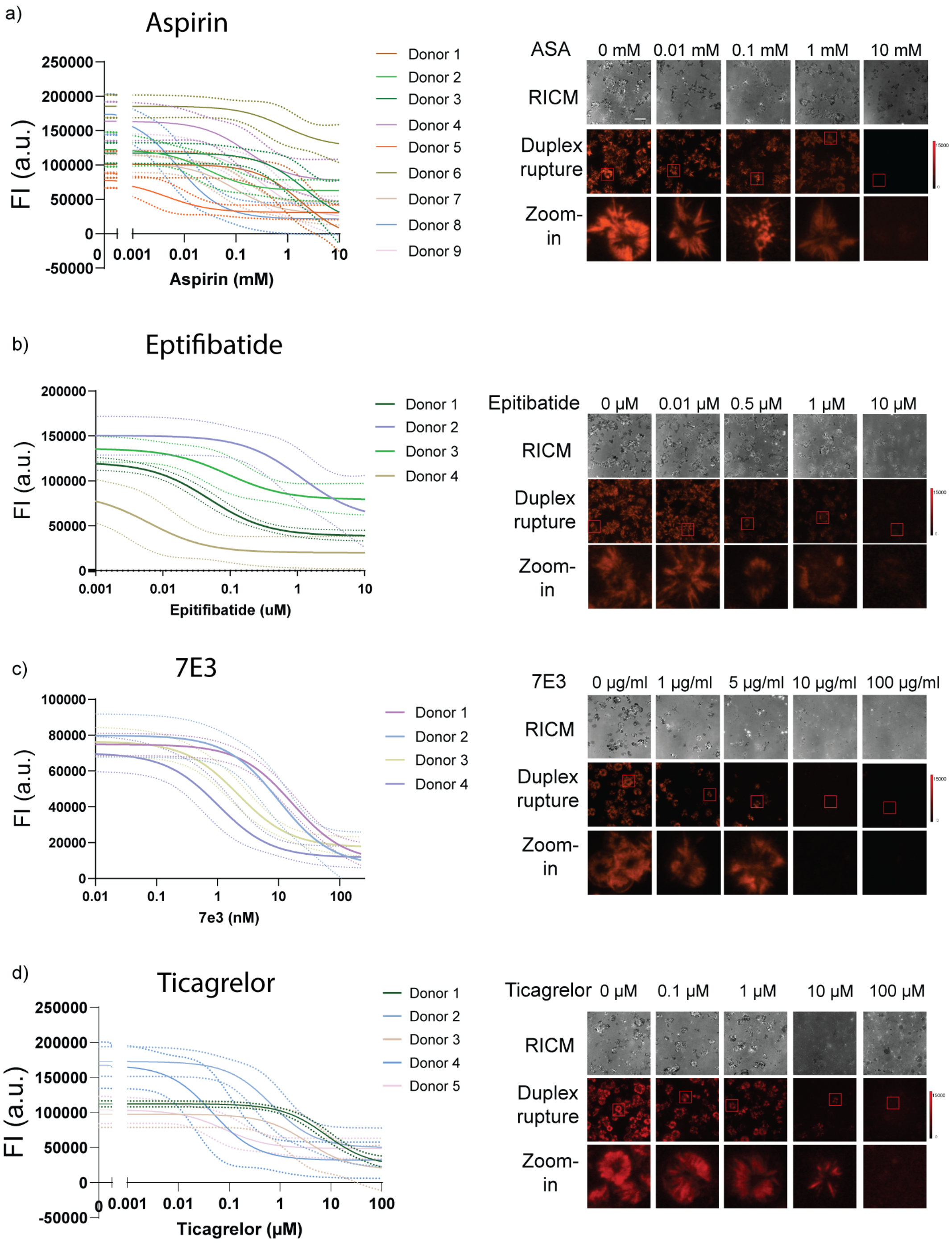

**Figure S10: MCATS used to measure dose-response curves for inhibitors of platelets.** a-d) Plots of [Aspirin], [Eptifibatide], [7E3], [Ticagrelor] vs MCATS signal for different donors. A dose-response titration of six drug concentrations for each drug for individual donors is measured with MCATS and all measurements were performed in duplicate or triplicate. Mechano-IC50 for each donor was calculated by fitting plot to a standard dose-response function: Signal=Bottom + (Top-Bottom)/(1+([drug]/IC50). Solid line represents the fitting and dashed line represents 95% CI. Representative RICM, duplex rupture fluorescence (red) and zoom-in fluorescence images for one donor with drug concentration ranging from 0 mM to 10mM. Scale bar = 10 μm.

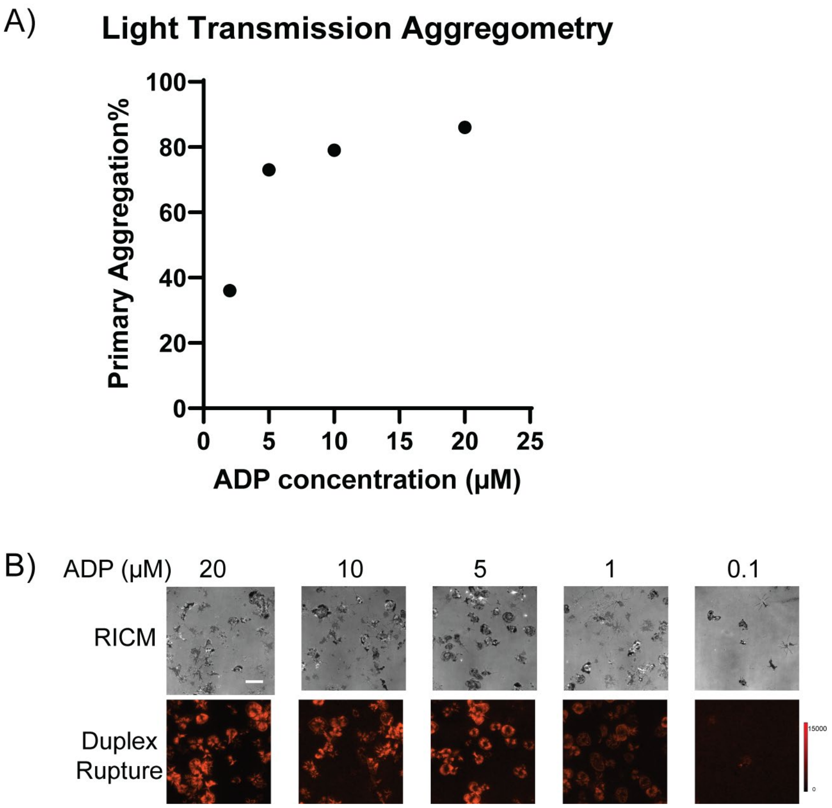

**Figure S11: LTA data for ADP agonist test.** A) Light Transmission aggregometry data for primary aggregation value against different concentration of ADP. B) Representative RICM, duplex rupture fluorescence (red) images for one donor with ADP concentration ranging from 0.1 mM to 20mM. Scale bar = 10 μm.

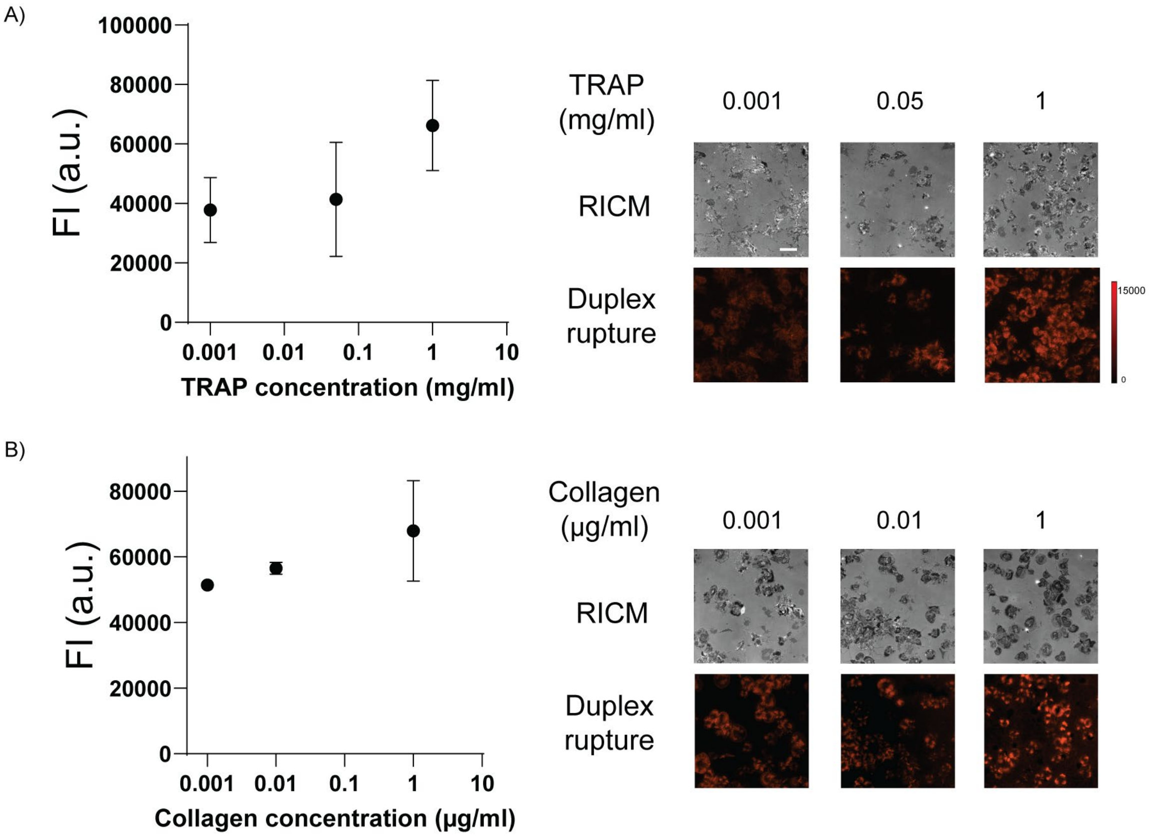

**Figure S12: TRAP and collagen agonist test.** A) Plots of MCATS signal against different concentration of TRAP. Error bar represents SD from n=2 measurements. Representative RICM, duplex rupture fluorescence (red) images for one donor with TRAP concentration ranging from 0.001 mg/ml to 1mg/ml. Scale bar = 10 μm. B) Plots of MCATS signal against different concentration of Collagen. Error bar represents SD from n=2 measurements. Representative RICM, duplex rupture fluorescence (red) images for one donor with TRAP concentration ranging from 0.001 μg/ml to 1μg/ml. Scale bar = 10 μm.

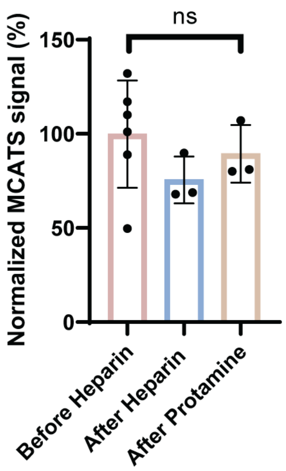

**Figure S13: Heparin and protamine influence on platelets tension.** Plots of normalized MCATS signal for purified platelets from non-heparin treated blood, purified platelets from 2hr heparin (0.25U/ml) treated blood, and purified platelets from heparin treated and protamine neutralized (1mg/100U) blood. Results indicating heparin influence on platelets was fully reversed after protamine addition. Error bar represents S.E.M. from n=3 independent experiments.

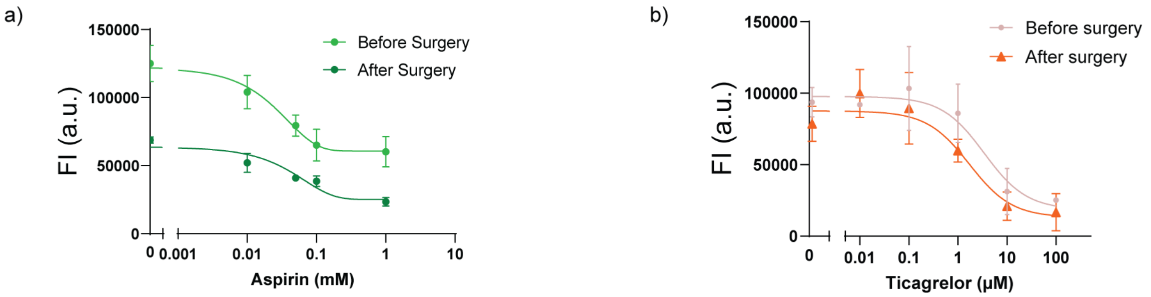

**Figure S14: Sensitivity of aspirin and ticagrelor for patients before and after surgery.** (A) Plots of [Aspirin] vs MCATS signal for patient before and after Surgery. Error Bar representative three replicate wells in the same day experiment. Mechano-IC50 for each donor was calculated by fitting plot to a standard dose-response function: Signal=Bottom + (Top-Bottom)/(1+([drug]/IC50). The difference of calculated $IC_{50}$ before and after surgery for two patients is non-significance with two tailed student T test, *P*=0.65. (B) Plots of [Ticagrelor] vs MCATS signal for patient before and after Surgery. Error Bar represent two replicate wells in the same day experiment. Mechano-IC50 was calculated by fitting plot to a standard dose-response function: Signal=Bottom + (Top-Bottom)/(1+([drug]/IC50).

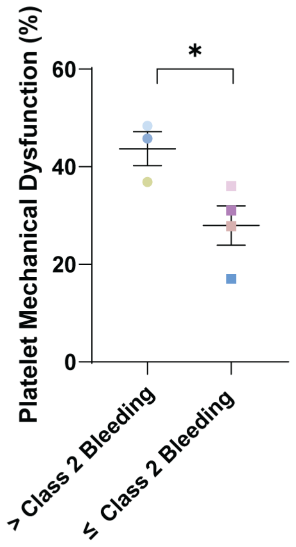

**Figure S15: Patient bleeding severity is correlated with platelet mechanical dysfunction.** Plot of reduction in MCATS signal (%) for subjects binned into two groups: mild/insignificant bleeding (class 1 and 2) or moderate/severe/massive bleeding (class 3, 4, and 5). Significance is calculated with two tailed student t-test with *P*=0.03. Error bars represent S.E.M.

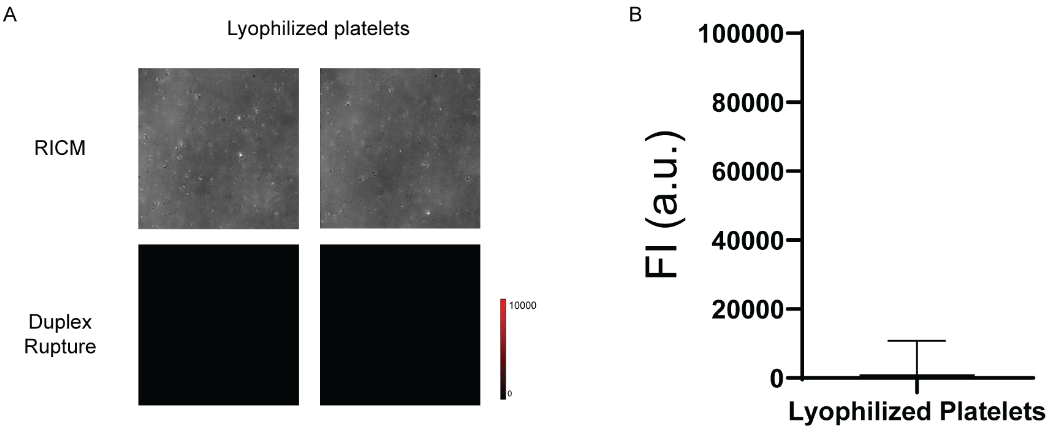

**Figure S16: Tension measurement of lyophilized platelets.** A) Representative RICM, duplex rupture fluorescence (red) images for lyophilized platelets on tension probes. B) Plots of MCATS signal of lyophilized platelets seeded on concealed DNA tension probes for 1hr. Results indicating lyophilized platelets have no active mechanical signal.

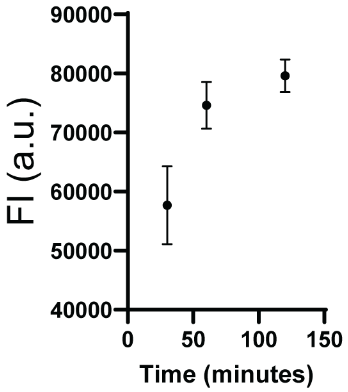

**Figure S17: Amplification time influence on MCATS signal.** Plots of MCATS signal at different time of amplification in platelet tension detection experiment. Error bar represents SEM of three independent experiments.

**Table S1: List of oligonucleotides**

| Name | Sequence (5' to 3') |
|---|---|
| **No Spacer Cas12a Bottom Strand** | /5AmMC6/CCTCCTCAAGGAACAACATTGC/3Bio/ |
| **Cas12a Top Strand** | /5Hexynyl/GCAATGTTGTTCCTTGAGGAGG/3BHQ_2/ |
| **6T Spacer Cas12a Bottom Strand** | /5AmMC6/CCTCCTCAAGGAACAACATTGCTTTTTT/3Bio/ |
| **60T Spacer Cas12a Bottom Strand** | /5AmMC6/GTGTCGTGCCTCCGTGCTGTGTTTTTTTTTTTTTTTTTTTT TTTTTTTTTTTTTTTTTTTTTTTTTTTTTTTTTTTTTTTT/3Bio/ |
| **60T Spacer Shearing mode Cas12a Bottom Strand** | /5BiosG/TTTTTTTTTTTTTTTTTTTTTTTTTTTTTTTTTTTTTTTTTTTT TTTTTTTTTTTTTTTT/iAmMC6T/CCTCCTCAAGGAACAACATTGC |
| **160T Spacer Cas12a Bottom Strand** | /5AmMC6/GTGTCGTGCCTCCGTGCTGTGTTTTTTTTTTTTTTTTTTTT TTTTTTTTTTTTTTTTTTTTTTTTTTTTTTTTTTTTTTTTTTTTTTTTTTTT TTTTTTTTTTTTTTTTTTTTTTTTTTTTTTTTTTTTTTTTTTTTTTTTTTTT TTTTTTTTTTTTTTTTTTTTTTTTTTTTTTTTTTTT/3Bio/ |
| **Control Scramble Bottom Strand** | CGCATCTGTGCGGTATTTCAC/3Bio/ |
| **Control Scramble top Strand** | GCTGGGCTACGTGGCGCTCTT |
| **ATTO 565N Reporter DNA** | /5ATTO565N/TTATT/3BHQ_2/ |
| **FAM reporter DNA** | /56-FAM/TTATT/3IABkFQ/ |
| **MCATS crRNA** | UAA UUU CUA CUA AGU GUA GAU GCA AUG UUG UUC CUU GAG GA |

**Table S2: ESI-Mass result**

| Strand Name | Expected Mass | Measured Mass |
|---|---|---|
| **Cy3B 60T Cas12a Bottom Strand** | 26059.7 | 26059.3 |
| **Atto 647N 60T Cas12a Bottom Strand** | 26145.7 | 26143.3 |
| **cRGDfk BHQ2 Cas12a Strand** | 8514 | 8514.7 |
| **Cy3B 6T Bottom Strand** | 9633.2 | 9632.8 |
| **Cy3B No spacer Cas12a Bottom Strand** | 7808 | 7807.5 |

**Table S3: Demographics, laboratory values, TEG data and surgery Note for CPB Patients**

| | Patient 1 | Patient 2 | Patient 3 | Patient 4 | Patient 5 |
|---|---|---|---|---|---|
| Age | 61 | 22 | 79 | 25 | 68 |
| Sex | F | M | F | F | M |
| Operation | MVR/TVR | PVR | Myectomy + AVR | Nephrectomy | Bentall |
| **Hematology*** | | | | | |
| Platelet count (cells mcL-1) | 322 | 264 | 172 | 482 | 168 |
| **Medication*** | | | | | |
| Aspirin | Y | Y | N | N | N |
| ADPI | N | N | N | N | N |
| **TEG6S MA (mm)** | 68.6 | 69 | 64 | 66 | 63 |
| **Operation Note** | | | | | |
| CPB time (min) | 314 | 84 | 73 | 204 | 182 |
| **Blood Products Transfused within 24h of Operation** | | | | | |
| RBC | 3 | 0 | 0 | 18 | 0 |
| PLT | 6 | 2 | 1 | 5 | 0 |
| FFP | 4 | 0 | 0 | 19 | 0 |
| Cryo | 15 | 5 | 0 | 50 | 0 |

| | Patient 6 | Patient 7 |
|---|---|---|
| Age | 71 | 39 |
| Sex | M | M |
| Operation | CABG x 3 | AVR/MVR |
| **Hematology*** | | |
| Platelet count (cells mcL-1) | 144 | 108 |
| **Medication*** | | |
| Aspirin | N | N |
| ADPI | N | N |
| **TEG6S MA (mm)** | 65.4 | 68.1 |
| **Operation Note** | | |
| CPB time (min) | 116 | 405 |
| **Blood Products Transfused within 24h of Operation** | | |
| RBC | 0 | 6 |
| PLT | 0 | 3 |
| FFP | 0 | 7 |

| | | |
|---|---|---|
| Cryo | 0 | 0 |
